# Accurate Drift-Invariant Single-Molecule Force Calibration Using the Hadamard Variance

**DOI:** 10.1101/2024.06.17.599270

**Authors:** Stefanie D. Pritzl, Alptuğ Ulugöl, Caroline Körösy, Laura Filion, Jan Lipfert

## Abstract

Single-molecule force spectroscopy (SMFS) techniques play a pivotal role in unraveling the mechanics and conformational transitions of biological macromolecules under external forces. Among these techniques, multiplexed magnetic tweezers (MTs) are particularly well suited to probe very small forces, ≤1 pN, critical for studying non-covalent interactions and regulatory conformational changes at the single-molecule level. However, to apply and measure such small forces, a reliable and accurate force calibration procedure is crucial.

Here, we introduce a new approach to calibrate MTs based on thermal motion using the Hadamard variance (HV). To test our method, we develop a bead-tether Brownian dynamics simulation that mimics our experimental system and compare the performance of the HV method against two established techniques: power spectral density (PSD) and Allan variance (AV) analyses. Our analysis includes an assessment of each method’s ability to mitigate common sources of additive noise, such as white and pink noise, as well as drift, which often complicate experimental data analysis. Our findings demonstrate that the HV method exhibits overall similar or even higher precision and accuracy, yielding lower force estimation errors across a wide range of signal-to-noise ratios (SNR) and drift speeds compared to the PSD and AV methods. Notably, the HV method remains robust against drift, maintaining consistent uncertainty levels across the entire studied SNR and drift speed spectrum. We also explore the HV method using experimental MT data, where we find overall smaller force estimation errors compared to PSD and AV approaches.

Overall, the HV method offers a robust method for achieving sub-pN resolution and precision in multiplexed MT measurements. Its potential extends to other SMFS techniques, presenting exciting opportunities for advancing our understanding of mechano-sensitivity and force generation in biological systems. Therefore, we provide a well-documented Python implementation of the HV method as an extension to the *Tweezepy* package.

**Statement of Significance:** Single-molecule force spectroscopy techniques are vital for studying the mechanics and conformations of bio-macromolecules under external forces. Multiplexed magnetic tweezers (MTs) excel in applying forces ≤ 1 pN, which are critical for examining non-covalent interactions and regulatory changes at the single-molecule level. Precise and reliable force calibration is essential for these measurements. In this study, we present a new force calibration method for multiplexed MTs using Hadamard variance (HV) based on thermal motion. The HV method shows similar or even higher precision and accuracy to established techniques like power spectral density and Allan variance. Most significantly, it is drift-invariant, maintaining consistent performance across varying experimental conditions. This robustness against drift ensures reliable force application and measurements at sub-pN resolution.

## Introduction

Single-molecule force spectroscopy (SMFS) techniques have emerged as powerful tools to investigate the behavior of biological macromolecules under forces and torques. ^1-6^ SMFS measurements have provided comprehensive insights into various biological processes, such as e.g. the function of molecular motors and DNA and protein mechanics^6-12^, interaction potentials between receptor-ligand pairs or DNA and different binding agents^13-15^, as well as protein folding pathways and associated dynamics^16-18^. For many biologically relevant questions, it is critical to resolve low forces in the sub-pN range (≤ 1 pN), e.g. to reveal specific protein interactions or small conformational changes involved in regulatory processes including cell motility, development, and differentiation. Here, magnetic tweezers (MTs) offer several important advantages compared to other established SMFS techniques such as optical tweezers and atomic force microscopy: ^5, 19-21^ MTs provide access to a large force range of ∼0.01-100 pN with high stability and accuracy and high-resolution in particular at low forces and in a multiplexed format to measure hundreds of molecules at the same time.^1, 6, 12, 22-26^ The latter is essential to obtain sufficient statistics from a single experiment, to observe rare events and identify molecular subpopulations, and to test diverse conditions and stimuli. Furthermore, MT measurements are intrinsically at constant force without a need for feedback control^12^ and do not suffer from heat generation or photo-damage^27^, which permits stable measurements on time scales ranging from sub-milliseconds to weeks.^22, 28^ Notably, various magnet geometries have been used to not only apply constant forces, but also to control and monitor torque and twist at the molecular level.^29-31^

In MTs, molecules of interest are tethered between a functionalized surface and small superparamagnetic beads (Figure 1A). External magnets exert controlled forces and torques on the beads and subsequently to the molecule of interest. To achieve highly parallel or multiplexed measurements, cameras with tens of mega-pixels can be used to image large fields of view (≥ 0.5 mm^2^). Hence, tracking of tens to hundreds of tethered beads at the same time becomes possible (Figure 1B), while applying constant forces across the field of view.^22^ A first key step in SMFS measurement is to perform an accurate force calibration in order to obtain exact and reproducible results. The most common approach for calibrating the forces is to make use of the bead’s thermal motion^32-34^. This method makes use of the equipartition relation that links the variance of thermal motion to the trap stiffness of the system and the thermal energy *k*_*B*_*T*, where *k*_*B*_ is the Boltzmann constant and *T* the temperature. Since the temperature of the system is easy to measure, calibration using thermal motion tends to be more accurate and reliable than alternative methods based on e.g. bead sedimentation dynamics^27, 35, 36^ or calculations of the magnetic field and exerted force^37^.

**Figure 1:**
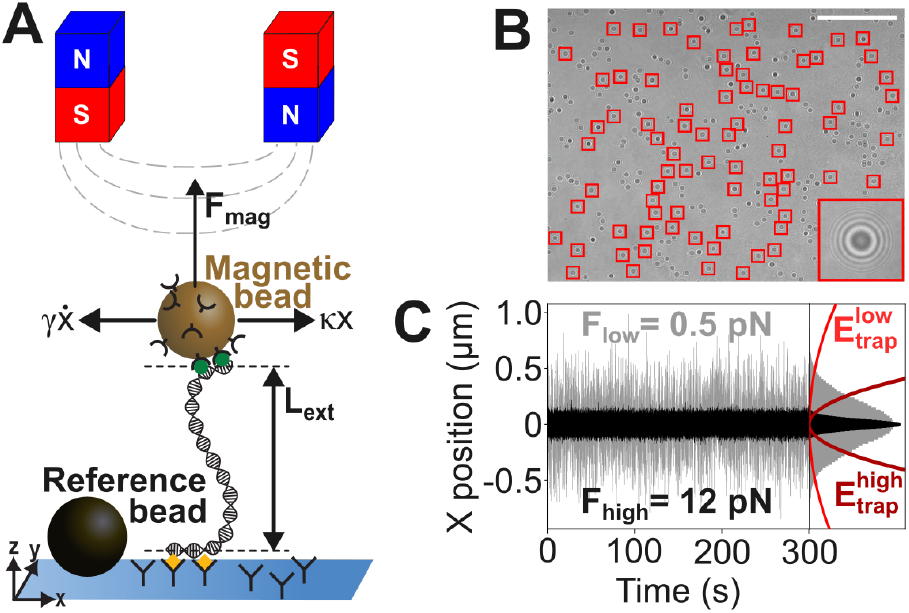
Schematic of a magnetic tweezers setup. **A**) The DNA with an extension *L*_*ext*_ is tethered between a substrate and a superparamagnetic bead. External magnets exert a magnetic force (*F*_*mag*_) on the bead. Drag *γ* and trap stiffness *κ* act on the bead parallel and vertical to the substrate in all three dimensions x, y and z (for simplicity, only highlighted in x direction). **B**) Field of view of our MT setup. Red squares highlight 78 beads that were tracked over time. The inset shows a zoom of one bead’s diffraction ring pattern used for tracking. The scale bar is 100 μm. **C**) Example (simulated) time traces of transverse fluctuations for different magnetic forces (left) and trap energy landscapes (right). The histograms on the right are computed from the time traces on the left.

In the thermal-motion based force calibration procedure, the bead is trapped in a harmonic potential originating from the combination of forces from the external magnetic field and from the elastic response of the molecular tether^34^. One calibrates the forces by analyzing the thermal trajectory of the bead.^32^ Typically, long (> 1 µm in contour length) dsDNA molecules are used for this force calibration method, due to their well-known properties and force-response characteristics. In particular, at low forces the force-extension relation for double-stranded DNA is well-described by the worm-like chain model.^38-40^ In addition, DNA undergoes a sharp overstretching transition at ∼ 65 pN resulting in a structural transition and corresponding abrupt length increase.^41^ In addition, long dsDNA molecules make it relatively easy to accurately determine the thermal motion of the attached magnetic beads for several resons:^42-44^ First, tracking errors (typically 1-2 nm) matter less for long tethers.^45-47^ Second, long DNA tethers result in larger characteristic times and hence slower thermal bead motion, which reduces the impact of finite camera exposure times.^34, 42, 48, 49^ Third, off-center attachment^50^ of a long DNA strand to the magnetic bead becomes less of an issue since the extension of the tether is much larger than the bead radius.

At long observation times (*τ* ≫ 10^−4^ s), hydrodynamic effects between the bead and its aqueous environment can be neglected^51^ and the bead motion is well-described by the overdamped Langevin equation, which contains two parameters, the drag coefficient *γ* of the bead and the harmonic trap stiffness *κ*.^42^ Using the equipartition theorem, the trap stiffness *κ*, which in turn is related to the force, can be determined from the transverse thermal motion of the tethered magnetic bead.^33, 34^ Specifically, for a magnetic bead confined to a harmonic trap, the equipartition theorem gives the real-space variance (RSV) by

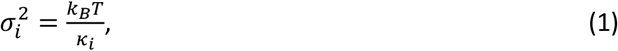

where *i* ∈ {*x, y, z*}, *k*_*B*_ is the Boltzmann constant, and *T* is the temperature. Here *κ*_*i*_ is the spring constant of the trap along the *i*^*th*^direction. Hence, stiff and soft traps result in small and large variances of the bead trajectory, respectively (Figure 1C). Conversely, if we measure the bead displacement at a known temperature *T*, we can calculate the trap stiffness *κ*_*i*_ for each direction.

In particular, the variance of bead motion in the transverse direction parallel to the magnetic field is directly related to the applied force *F*_*z*_ via^34^

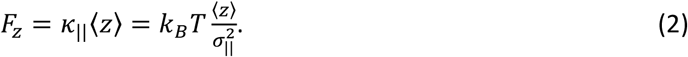

where ⟨*z*⟩ is the mean extension of the molecule in the *z* direction. Note that in principle, we could make use of the fluctuations which are perpendicular to both the *F*_*z*_ and the magnetic field, but in that case the measured extension of the molecule must be corrected to include rotation of the bead in the magnetic field.^42^

While conceptually straight forward, the RSV approach has two key limitations: First, due to the finite shutter time in camera-based tracking, the acquired signal is distorted due to motion blurring and temporal aliasing (termed downsampling throughout this work, Figure 2A), which act as a low-pass filter and lead to under-estimation of the true variance.^42, 48, 49^ Hence, spatiotemporal information is lost if the sampling frequency is too low, which leads to an over-estimation of the applied forces (Figure 2B). Errors due to the finite shutter time are particularly problematic for short tethers and high forces since the characteristic time of the intrinsic fluctuations decreases with increasing force and decreasing tether length. A number of approaches have been put forward to overcome the limitations due to finite shutter time, including high-frequency tracking^43, 47^, shuttered illumination^44, 48^, and the application of spectral corrections in the analysis^42^.

**Figure 2:**
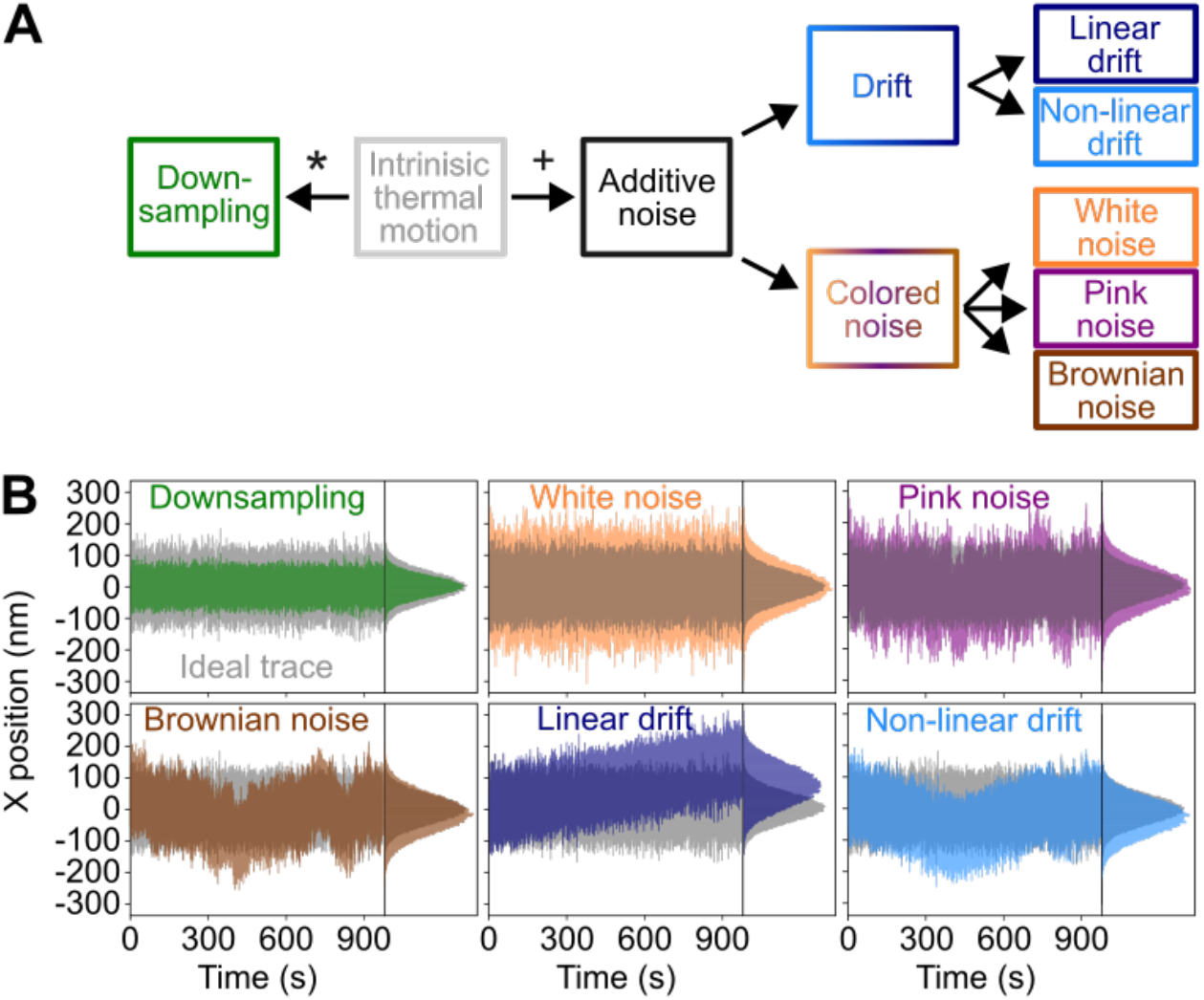
Overview of processes affecting time traces of tracked beads in MTs. (**A**) Classification of the spatial and temporal alterations of the intrinsic thermal motion. Downsampling acts as a low pass filter introducing blurring and aliasing and is denoted as “ * “, as it effectively acts as a convolution of the signal. Additive noise contributions denoted as “ + “. We differentiate between drift (linear and non-linear) and colored noises (white, pink, Brownian noise). (**B**) Example traces for downsampling and different additive noise types. The unaltered (ideal) thermal motion bead trace for a sampling frequency of 10 kHz is shown in grey in all panels; the same trace downsampled to 72 Hz is shown in green. White, pink, and Brownian noise, and linear and non-linear drift contaminated trace for signal-to-noise ratios of –2.4 dB, –1.6 dB, 0.6 dB, –9.7 dB and 0.9 dB, respectively, are shown in the colors indicated in the legend. All traces were obtained via Brownian dynamics simulations (see Methods) assuming a force of 2.5 pN and a 1 μm bead. The individual SNRs were chosen for visualization purposes.

The second key limitation of the RSV approach is that it is susceptible to any kind of additive noise^52^, which affects magnetic tweezers measurements by introducing spatial alterations and bias to fitting algorithms^53^ (Figure 2B). Note that we explicitly distinguish here between the intrinsic thermal Brownian bead motion (sometimes also referred to as “noise” in the literature), which is our “signal” in the context of force calibration, and additional detrimental spatial alterations, which we call “additive noises” (Figure 2A). Among the types of additive noises, colored noises, which follow a power-law spectrum, are well-known sources for spatial distortions. There are three common types of colored noise that can, in particular, cause erroneous force calibration of SMFS systems (Figure 2B): First, the use of electric devices such as cameras in MT settings for data acquisition and computer-based signal processing generates white noise.^54, 55^ White noise is an additive type of colored noise resulting in broadening of the bead trajectories and thereby under-estimation of the force. Second, resistance fluctuations in the electronic components (camera, motors, etc.)^54^ and structural relaxations of the DNA^56, 57^ (overstretching, solvent effects, etc.) can induce pink noise. Pink noise is also known as flicker noise and leads to random spatial shifts of the bead trajectory. Third, similar spatial shifts can also emerge from unaccounted physical processes (light-matter interactions, rotational bead motion, etc.) that generate Brownian noise^58^ in addition to the thermal bead motion one wants to extract and analyze. Like white and pink noise, Brownian noise broadens the bead trajectories spatially and causes force under-estimation.

To address the challenges associated with downsampling and colored noises, a number of extensions of Eq. 1 have been introduced to obtain a better estimation of *κ*‖. These calibration methods include (1) the power spectral density (PSD) and iterative corrections of the camera effects^42^, (2) the PSD with additional correction factors for tracking errors and colored noises^49^, and (3) the Allan variance^49^ method which accounts for frequency instabilities. However, none of these calibration methods can handle drift, which is also a type of additive noise, and a well-known issue in SMFS experiments, in addition to downsampling and colored noises. While mechanical drift is commonly corrected for in MTs by tracking reference beads^52^, residual, uncorrected drift still remains in the data.^46^ In particular in multiplexed MTs with a large field-of-view, drift across the field-of-view can be an additional limiting issue (Figure 2B). The Hadamard variance (HV)^59^ has been shown to offer significant advantages in handling drift in metrology contexts, as it uses a second-order finite difference method that renders it drift-invariant by definition by cancelling linear contributions.^60^ Hence, in theory, the HV is more robust with respect to drift than the PSD and AV methods. Furthermore, compared to the AV, the HV exhibits higher resolution in estimating the frequency spectrum since it has a smaller noise-bandwidth.^61^

To extend the existing force calibration methods for multiplexed MT measurements, here we present a thermal motion-based model of the Hadamard variance (HV). We compare all three methods, i.e. PSD, AV and HV, using simulated and experimental data traces. First, we test our method with simulated bead trajectories at forces ranging from 0.5 pN to 80 pN. We compare the thermal motion of an ideal trace to traces with various sources of error, including downsampling, white noise, pink noise, Brownian noise, and drift, at experimentally relevant signal-to-noise ratios (SNRs) and drift speeds. We simulate data with SNRs of −10 to +30 dB, which correspond to 10 times higher noise compared to signal levels, and 1000 times higher signal compared to noise levels, respectively. Overall, we make two important observations: First, all three methods can account for white, pink, and Brownian noise with the HV method being more precise at higher SNRs of ≥ 10 dB. Second, the HV methods is the most stable and accurate in presence of linear and non-linear drift over the whole studied signal-to-noise and drift speed ranges. Finally, application of the methods to experimental data and quantification of the goodness-of-fit show that the HV method gives the most precise force estimation values, as we find the relative force errors for the HV to be lower compared to PSD and AV. Overall, our findings highlight that the Hadamard variance is a suitable tool to process multiplexed single-molecule force spectroscopy data sets to obtain accurate and precise results in the presence of colored noise and drift, which we expect to be useful for the calibration of force spectroscopy techniques and related applications.

## Methods and Materials

In order to determine the accuracy of the PSD, AV, and HV methods in handling downsampling, colored noise and drift, we employ simulations, analytical theory, and experimental measurements. Here, we first describe our methodology to simulate the Brownian dynamics of the bead-tether system, including downsampling, colored noise, as well as linear and non-linear drift. Second, we summarize the theoretical background and formulae for the PSD, AV, and HV methods, and third, the experimental details of the MT setup and force calibration measurement are reported.

### Simulations of the bead-tether system and inclusion of downsampling, colored noise, and drift

#### Brownian-motion simulations

Our goal is to mimic an experimental MT setting where a long dsDNA tether (21 kbp) is tethered between a bead and a surface. We model this as a bead-tether system in three dimensions (3D). We assume that the bead is pulled by a constant magnetic force perpendicular to the hard wall and we model the DNA tether as a worm-like-chain (WLC) whose contour (*L*_*c*_) and persistence (*L*_*p*_) lengths are 7 μm (corresponding to 21 kbp) and 45 nm, respectively^62^. Furthermore, collisions between the magnetic bead and solvent molecules are included as a 3D stochastic Langevin force ***F***_***L***_ that depends on the proximity to the hard wall due to hydrodynamic effects. The 3D stochastic Langevin force ***F***_***L***_ obeys the fluctuation-dissipation theorem 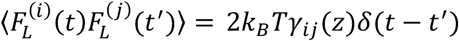 with *δ*(*t* − *t*′) being the Dirac distribution and *γ*_*ij*_(*z*) the Faxén-corrected^62-64^ diagonal viscous drag matrix elements that account for the hydrodynamics of the hard wall on the bead motion. By combining these effects, we arrive at the overdamped Langevin equation that describes the motion of the magnetic bead as

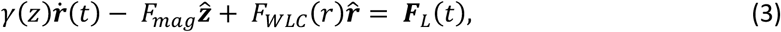

with ***r*** = (*x, y, z*) being the position vector 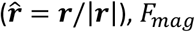 the magnitude of the magnetic force, and *F*_*WLC*_(*r*) the WLC force given by the 7-parameter WLC model^40^.

To numerically solve the equation of motion, we write Eq. (3) in the stochastic differential equation (SDE) form,

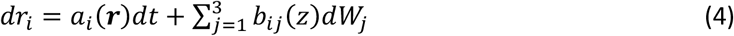

where 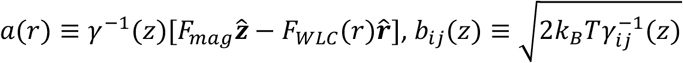 and *W*_*j*_ are independent Wiener processes. Then, we employ the Milstein method^65^ to cast the SDE into the recursive relation,

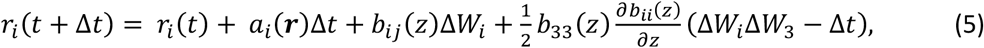

where Δ*W*_*i*_∼*N*(0, Δ*t*) is a Gaussian distributed random number with zero mean and variance Δ*t*. The custom Python 3 code implementing our Langevin simulations of the bead-tether system is available at XXX.

Using the simulation method introduced above, we perform simulations at various magnetic forces (0.5 pN, 2.5 pN, 12.0 pN, 40.0 pN, 80.0 pN) and two different micro-sphere sizes, representing the commonly used MyOne (diameter: 1 μm) and M270 (diameter: 2.8 μm) superparamagnetic beads.

#### Downsampling of simulation trajectories

We include downsampling as follows: We divide the simulation trajectory into non-overlapping windows of width 1/*f*_*s*_ *with f*_*s*_ *being the sampling frequency*. Then, we calculate the average of each window, yielding samples separated by 1/*f*_*s*_ that form the desired downsampled trajectory.

#### Addition of colored noise and drif t to simulation trajectories

To add colored noises, we follow a four step recipe^66^. (1) First, we generate independent sets of Gaussian random numbers whose length matches the trajectory length, which corresponds to white noise. (2) Then, we take the Fourier transform (FT) of the noise and rescale the frequency contributions to match the power spectral density profile of the colored noise, followed by taking an inverse FT to revert to real space. (3) Next, we calculate the powers of the pure signal and the noise by numerically integrating their autocorrelation function and we rescale the noise to match the desired (−10 to +30 dB) signal-to-noise ratios (SNR). (4) Finally, we add the rescaled noise to the ideal signal to obtain the noisy trace with a defined noise type and magnitude. To add linear drift to the trajectory, we add a line with defined slope, i.e. drift speeds of 0-50 nm·s^−1^. To add non-linear drift, we add colored noise with a power spectral density that is strongly peaked at low frequencies (*S*(*f*) ∝ 1/*f*^4^), using the same procedure as for the other colored noises.

### Closed-form expressions for the power spectral density, Allan variance, and Hadamard variance

In order to use the PSD, AV, or HV to determine *κ*, we require expressions that relate measurements of these quantities to *κ*. To this end, we consider a simple model of the bead-tether system in one dimension. Specifically, we model the whole bead-tether system as a harmonic trap of stiffness *κ* centered at the equilibrium position. Collisions between bead and solvent molecules are modeled by a stochastic Langevin force *F*_*L*_ that obeys the fluctuation-dissipation relation given as ⟨*F*_*L*_(*t*)*F*_*L*_(*t*′)⟩ = 2*γk*_*B*_*Tδ*(*t* − *t*′) with *γ* being the viscous drag coefficient. The bead’s motion is then well-described by the overdamped Langevin equation

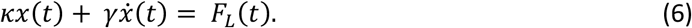

One way to infer the trap stiffness *κ* is the use of the RSV method, as described in the introduction. However, to also account for downsampling and colored noise types including, white, pink, and Brownian noise, closed-form expressions of the PSD and AV have been derived for the overdamped Langevin equation (Eq. (6)) in order to extract the trap stiffness *κ* (see below). While the PSD and AV approaches have been used widely to obtain force estimates from tweezer time traces, they can be susceptible to drift and other measurement imperfections. Therefore, we derive here a closed-form expression of the Hadamard variance for Eq. 6., to establish an accurate drift-invariant force calibration technique.

#### Power spectral density

The PSD is a measure of the power contribution of the different signal frequency components to the total signal given as

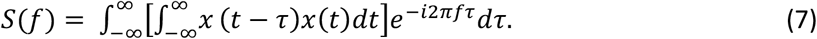

The PSD *S*(*f*) of the bead trajectory then follows from Fourier analysis of the overdamped Langevin equation (Eq (6)). The downsampling-corrected analytical closed-form expression of the PSD is given by _49, 67_

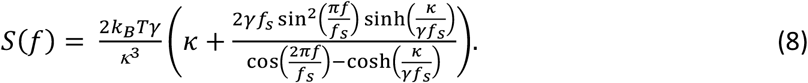

with *f*_*s*_ the sampling frequency. Notably, Eq. (8) holds for the special case of equal finite camera exposure time *τ*_0_ and sampling time *τ*_*s*_ = 1/*f*_*s*_, i.e. *τ*_0_ = *τ*_*s*_. ^67^ This assumption is applicable to most modern video cameras because the camera dead time (∼10^−6^ s) is in general much smaller than the sampling time (∼10^−4^ s to 10^−1^ s). The PSD has a Lorentzian-like form with a turning point at the corner frequency (Figure 3A).

**Figure 3:**
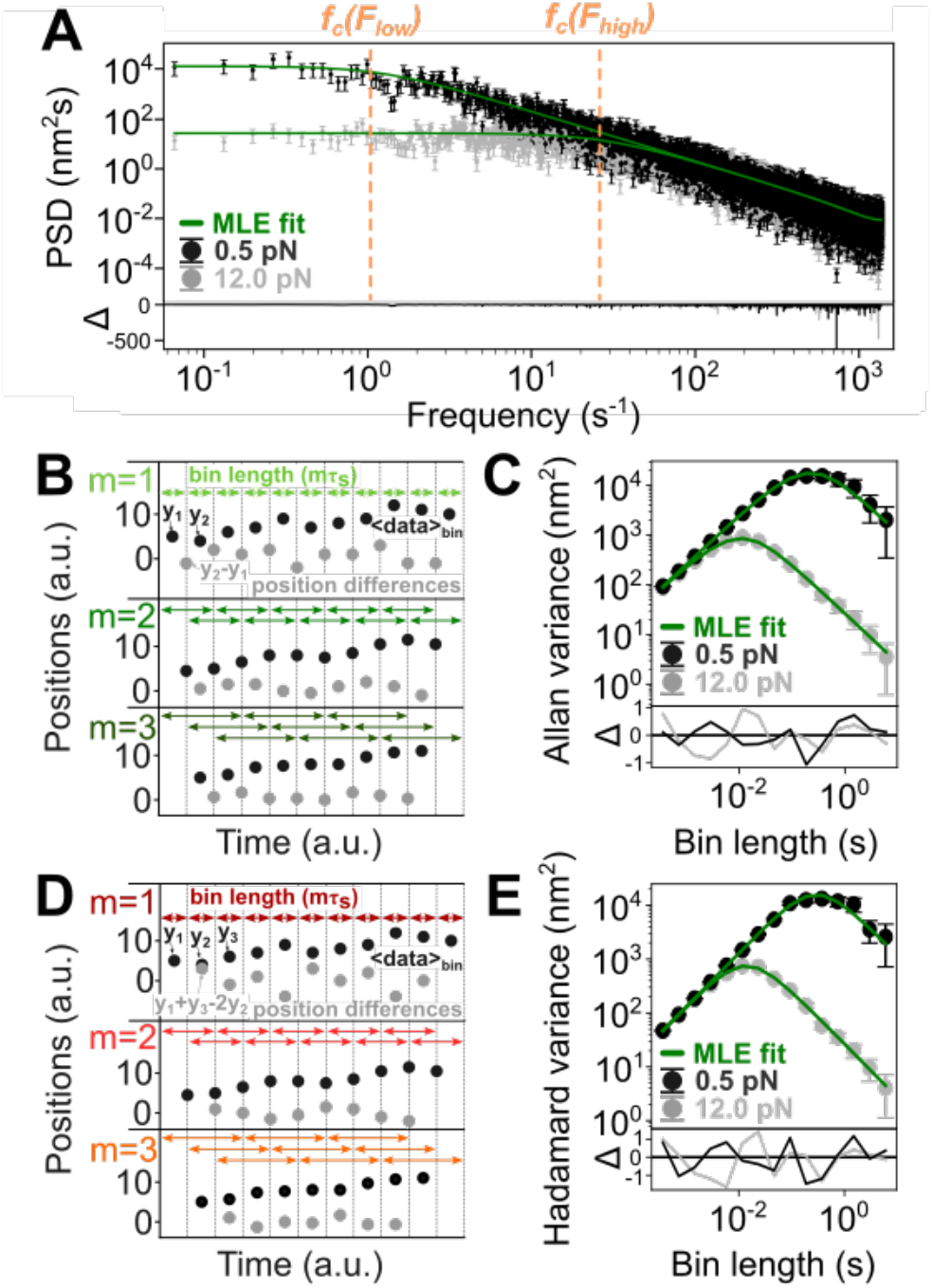
Illustration of the PSD, AV, and HV analyses. Example plots of the PSD (**A**), Allan variance (**C**) and Hadamard variance (**E**) methods applied to ideal, simulated bead trajectories. The lower parts (Δ) represent the residuals of the MLE fit to the simulated data. **(B, D)** Schematic of the position difference calculations for the Allan (B) and Hadamard (D) variance. The black dots for m = 1 are the original signal. The black dots for m = 2 and 3 are the position signals averaged over bin comprising m points. The (first and second order) position differences are shown as grey dots and are derived from two and three consecutive data points for Allan and Hadamard variance, respectively. The arrows represent the bin lengths.

#### Allan variance

The AV of the bead’s thermal motion is computed by separating the signal into different observation time intervals or bins of length *τ*, and calculating the first position differences (first-order finite difference method) i.e. the differences of two consecutive averaged data points, as a function of the observation time bins (Figure 3B,C). Specifically, we calculate the Allan variance from the bead trajectories via^68^

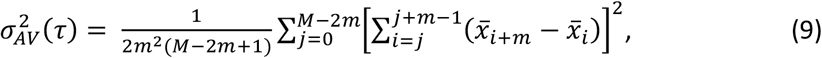

with 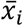 the position average of *i*^*th*^ bin. The position averages 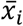 of the positions *x*_*n*_ are given by

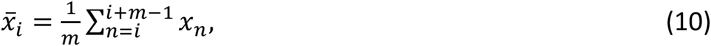

with *m* = *τ*/*τ*_*s*_ (*τ*_*s*_: sampling time) the overlapping bin lengths of the observation time intervals. The bin lengths define the number of bins *M* = *N* − 2*m* + 1 with *N* the total number of data points. Note that this corresponds to the so-called overlapping Allan variance due to the consideration of overlapping time intervals.

The closed-form expression of the AV for Eq. (6) is then given by ^49^

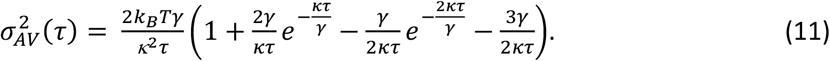

Hence, in order to use the AV to obtain *κ*, we calculate the AV of the measured x trajectory of the bead. Then, we follow a strategy reported by Morgan *et al*. ^67^ to fit the calculated AV to Eq. (11) with *κ* and *γ* being fitting parameters using maximum likelihood estimation (MLE).

#### Hadamard variance

In contrast to the AV, the HV is based on the calculation of the second position differences (second-order finite difference method), which eliminates any additive linear signal contribution and thus renders it inherently insusceptible to linear drift. The overlapping Hadamard variance is defined as ^69^

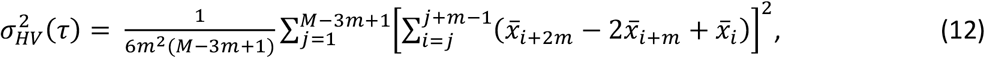

with 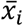 the position averages (Eq. 10, Figure 3D). The number of bins of the overlapping HV is given as *M* = *N* − 3*m* + 1.

The closed-form expression of the HV of the bead motion (Eq. (6)) then follows as

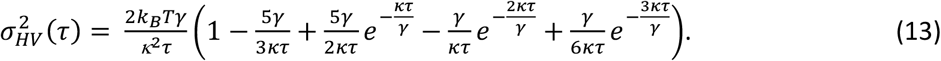

The full derivation of the expression in Eq. 13 is given in the Supporting Information, Section S1. To implement a computational method to calculate the HV from the bead trajectories and determine the trap stiffness and drag coefficient from the HV curves, we followed a similar strategy as Morgan *et al*. ^67^ employing an MLE fit of the HV as a function of observation times (Figure 3E). We implemented the HV method in a python framework that extends the *Tweezepy* package^67^ that can be downloaded from XXX.

Note that in principle it is possible to include spectral correction factors for white, pink, and Brownian noise in Eqs. 8, 11, and 13, allowing one to correct for such types of noise when they are well understood and quantified. However, as the purpose of this study is the quantification of each methods’ susceptibility to these types of additive noises, we do not include these corrections here.

### Magnetic tweezer experiments

#### Magnetic tweezer setup

We use a custom build MT setup equipped with two linear stepper motors (Physik Instrumente, Germany) to control the height of an external magnet (1 mm gap) above the flow cell and its rotation, and a Pifoc motor (Physik Instrumente, Germany) to adjust the focus of the objective (40x oil immersion, Olympus, Japan). ^17, 37, 70, 71^ Imaging of the diffraction rings of the magnetic beads uses a tube lens (f = 150 mm), a 650 nm LED (Thorlabs, Germany) for illumination, and an Optronis camera (CP80-25-M-72, Optronis, Germany) operated at sampling frequencies of 72 Hz (full field of view) or 400 Hz (reduced field of view). Bead tracking is achieved with an open source Labview code that is based on a CUDA parallel computing algorithm and quadrant interpolation^24, 45^.

#### DNA and flow cell preparation

We use a 21 kbp long dsDNA construct derived from lambda phage DNA, which was prepared as previously described^72^. The ends (∼600 bp) are functionalized for attachment with multiple digoxygenin (Roche, The Netherlands) and biotin (Sigma Aldrich, USA) molecules, respectively. Flow cells are built from two glass cover slips. The bottom slide (24 x 60 mm^2^ (#1), VWR, Germany) is first functionalized using (3-Glycidoxypropyl)-trimethoxysilane (Thermo Fisher Scientific, USA), then incubated with polystyrene beads (Polysciences, USA) of 1 or 3 µm diameter in ethanol (VWR, USA), which serve as reference beads for drift correction. The top slide (24 x 60 mm^2^ (#1), Carl Roth, Germany) contains two holes with diameters of ∼1 mm to connect the pump system (Ismatec, Germany) for fluid exchange in the flow chamber. Both slides sandwich a single layer of melted parafilm (Carl Roth, Germany) with a cutout to form a ∼50 µl channel that connects inlet and outlet of the flow chamber. After assembly, the flow cell is incubated with 200 µg/ml anti-digoxygenin (abcam, UK) in 1x PBS buffer (Sigma Aldrich, USA) for at least 2 h to allow later DNA attachment *via* digoxygenin-anti-digoxygenin binding. The channel is rinsed with 1x PBS and passivated with 100 mg/ml BSA (Carl Roth, Germany) for 1 h to avoid non-specific interactions, then rinsed again. We use commercial streptavidin-coated magnetic beads with diameters of 1 µm (DynaBeads MyOne Streptavidin C1, Thermo Fisher Scientific, USA) and 2.8 µm (DynaBeads M-270 Streptavidin, Thermo Fisher Scientific, USA) that are attached to the DNA *via* streptavidin-biotin binding. After coupling 1 µl of picomolar DNA stock with 2 µl MyOne beads or 13 µl M270 beads in 100 µl 1x PBS for around 30 s, we add the mixture to the flow cell and incubate for several minutes before flushing out unbound material with 1x PBS. The flow cell is mounted in a custom-built flow cell holder.

#### Force calibration measurements

For the force calibration measurements, we calibrate the distance between the magnet and upper cover slide of the flow cell by slowly approaching the top surface (to within 0.1 mm). We define the magnet position origin as the point where it touches the top coverslip. Next, the tethered magnetic beads are screened for attachment *via* multiple DNA molecules by measuring their response to force and torque. First, we apply negative turns (−70 turns) at intermediate forces (2 pN or 14 pN for MyOne or M270 beads, respectively), where no change in extension is expected for beads tethered by single double-stranded DNA molecules. Beads attached *via* multiple tethers form braids during negative rotation and thus decrease the extension. Multiple tethered beads are excluded from the measurements. Further, the tethers are screened for single strand breaks, by overwinding the DNA at low force. We apply positive turns (70 turns) at low forces (0.05 pN or 0.14 pN for MyOne or M270 beads, respectively) and record the tether extension. Nicked tethers are not able to form plectonemes when overwound under low force and thus no change in extension is expected. For the force calibration measurement, we record time traces of the bead-tether complex at 49 different magnet positions. We further record test traces at a low and high force i.e. a fully relaxed and an unconstrained stretched DNA tether, to set the tether-bead coordinate system. In addition, we identify off-center attached beads, which do not follow a 2D Gaussian position distribution in the x-y-plane using the Shapiro-Wilk test for normality (exclusion criterion: p-value < 0.05) and exclude them from the experiment.

## Results

We present and critically test different methods to analyze MT extension time traces for force calibration. To quantify and compare the accuracy of the PSD, AV, and HV methods in the presence of downsampling, colored noise, and drift, we first analyze simulated bead trajectories created by our Brownian dynamics simulations.

### Testing the PSD, AV, and HV analyses using simulated bead trajectories

As a first step, we simulate ideal (noise-free) thermal motion trajectories (Figure 2B and Materials and Methods). We choose simulation time steps that are two orders of magnitude smaller than the characteristic (corner) time and within the stability region of the Milstein method^73^. The corner frequency represents the turning point between purely diffusive bead motion and being constrained by the trap. To mimic the finite exposure time of the camera, we then downsample the trajectories by partitioning them into non-overlapping windows whose widths match the exposure time of the camera and taking the average of each window. Next, we add either white, pink, or Brownian noise, or linear or non-linear drift to the ideal thermal motion trace, since these are the most-common noise contributions to experimental MT data (Supporting Information, Section S2). To test the effects of different levels of additive noise, we apply noise with different signal-to-noise ratios (SNR). The SNRs (*SNR*_*dB*_ in dB) are calculated as the ratio of the power of the ideal thermal motion trace (*P*_*sig*_) to the powers of the pure additive noise (AN) traces *P*_*AN*_, i.e *SNR*_*dB*_ = 10 *log*_10_ (*P*_*sig*_/*P*_*AN*_). For linear drift, we add a linear trend of defined drift speed, or slope, to the ideal thermal motion trace.

To compare the three methods, i.e. PSD, AV, and HV, regarding their accuracy and precision in handling the different trajectory distortions, we next compute the PSD, AV, and HV for our simulated traces and fit the corresponding models, Equations 8, 11, and 13, respectively. We start by looking at 1 μm beads and two forces, 0.5 pN and 12 pN, and calculate the relative force errors according to |1 – (*F*_*method*_/*F*_*true*_)| with *method* comprising PSD, AV or HV (Figure 4). In the presence of downsampling, all three methods (PSD, AV, and HV) reach a relative error of ≈ 7 · 10^−2^ and ≈ 2 · 10^−2^ for 0.5 pN and 12.0 pN, respectively, at sampling frequencies that exceed twice the corner frequency, i.e. the Nyquist frequency (Figure 4A and Supporting Information, Section S3, Figure S2). A stable plateau of low relative force errors is thus reached at sampling frequencies that fulfill the Nyquist theorem, i.e. exceed twice the characteristic frequency of the fluctuations. We calculate Nyquist frequencies of 2.75 Hz and 54.81 Hz for 0.5 pN and 12 pN, respectively. At frequencies around and smaller than the Nyquist frequency, the relative force errors become on the same order as the applied force indicating that the methods are not invariant to downsampling, as expected from the Nyquist theorem. However, taking a closer look at the critical regime of sampling at the Nyquist frequency (*f*_*s*_ = *f*_*Nyquist*_), the HV method results in slightly lower force estimation errors compared to the PSD and AV methods for 0.5 pN and 12.0 pN (Figure 4B). Considering the error bars which correspond to one standard error, the differences between the methods are, however, statistically insignificant. Moreover, the PSD, AV, and HV methods achieve similar correction efficiencies, and all three methods perform significantly better than the RSV method (Supporting Information, Section S4).

**Figure 4:**
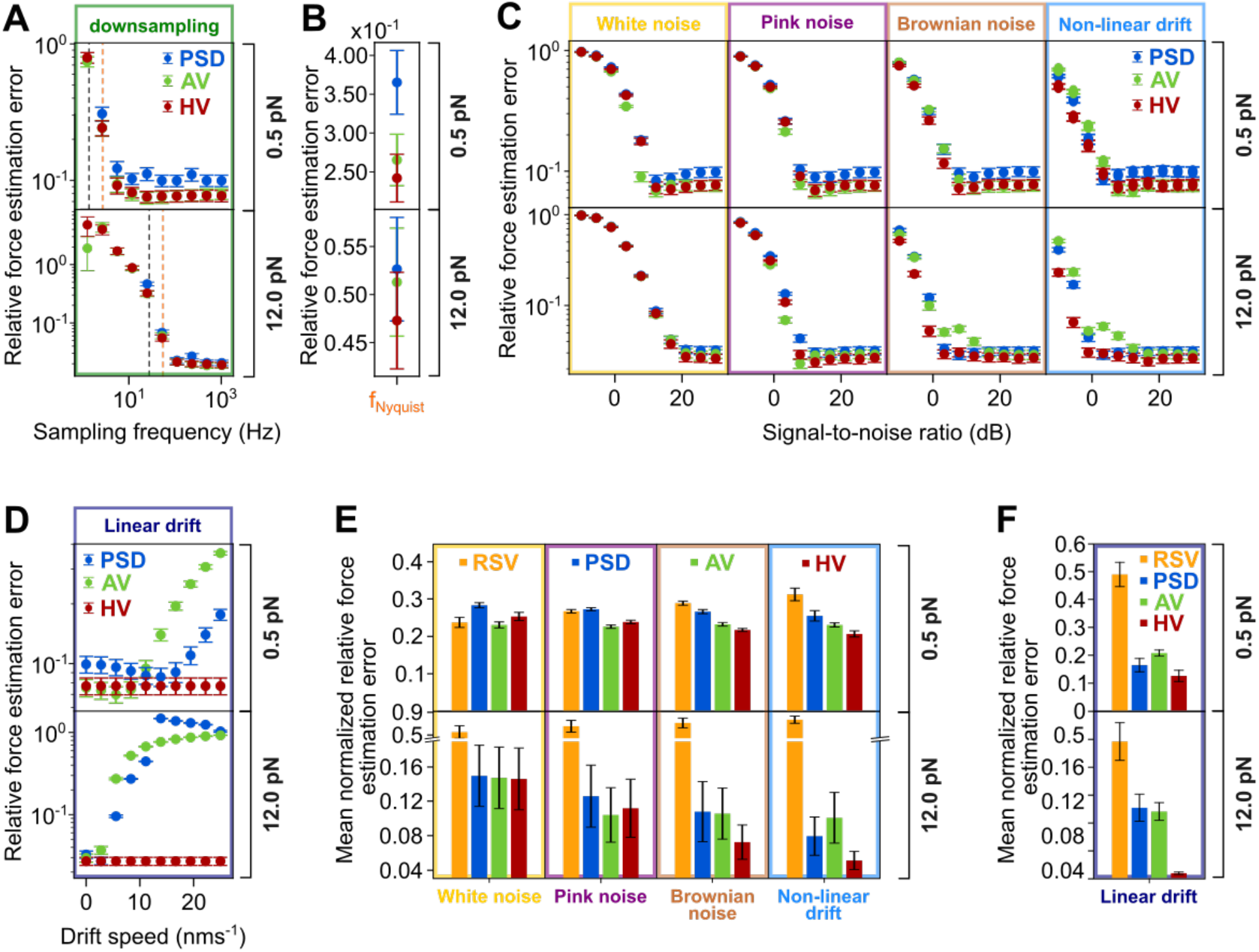
Effects of downsampling, colored noise, and drift on force estimation errors evaluated using simulated data. **A**) Relative force estimation errors as a function of the sampling frequency for PSD (blue), AV (green) and HV (red) at low (0.5 pN) and high (12 pN) force. The black dashed lines highlight the corner frequencies and the orange dashed lines the Nyquist frequencies. *Note:* The PSD method failed to convergence at frequencies smaller than the corner frequencies. **B**) Relative force estimation errors at the Nyquist frequency for PSD (blue), AV (green) and HV (red) at low (0.5 pN) and high (12 pN) force. **C)** Relative force estimation errors as a function of the signal-to-noise ratios for PSD (blue), AV (green), and HV (red) at low (0.5 pN) and high (12 pN) force (sampling frequency: 72Hz). **D)** Relative force estimation errors as a function of the drift speed for PSD (blue), AV (green), and HV (red) at low (0.5 pN) and high (12 pN) force (sampling frequency: 72Hz). **E**) Mean (SNR) normalized relative force estimation errors of the real-space variance (RSV, orange), PSD (blue), AV (green), and AV (red) at low (0.5 pN) and high (12 pN) force. The error bars represent the standard error of the mean. **F**) Mean (drift speed) normalized relative force estimation errors of the real-space variance (RSV, orange), PSD (blue), AV (green), and AV (red) at low (0.5 pN) and high (12 pN) force. The error bars represent the standard error of the mean.

For the traces containing colored noise, we find that the relative errors for PSD, AV and HV decrease with increasing signal-to-noise ratios in presence of white, pink, and Brownian noise (Figure 4C and Supporting Information, Section S5, Figures S4-S7), until the relative errors essentially plateau for high SNRs. We find that the HV is on average most accurate in handling colored noise at high signal-to-noise ratios ≥ 10 dB. At SNRs smaller than 10 dB no clear trend between the different methods can be identified. For trajectories containing non-linear drift, the HV clearly gives smaller force estimation errors over the whole SNR range compared to the PSD and AV methods.

Importantly, in presence of linear drift, the relative force estimation errors increase with drift speed for the PSD and AV methods but stay constant for the HV method, which highlights its robustness against linear drift (Figure 4D and Supporting Information, Section S6, Figure S8).

In addition, we also simulate traces of the 1 μm beads at an intermediate force of 2.5 pN and of the 3 μm beads (representing M270) over a force range of 0.5 pN to 80 pN and observe similar trends (Supporting Information, Sections S5 and S6).

To further give a more general overview of the force estimation accuracies of all methods, we next calculate the mean normalized relative force errors 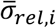 by normalizing the relative force estimation errors *σ*_*rel*_ with respect to the methods (*i, j*) and summing over all SNRs or drift speeds using 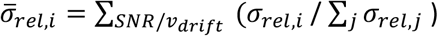 (Figures 4D,E). For comparison, here we also include the real-space variance (RSV) method described in the introduction (Eq. 1). Note that a full analysis of the RSV in presence of downsampling and additive noises is included in the Supporting Information, Section S7.

As illustrated in Figure 4D, all four methods, i.e RSV, PSD, AV, and HV, achieve almost equal accuracy with similar mean normalized force estimation errors at low force (0.5 pN) and in presence of white and pink noise. However, the good apparent performance of the RSV is misleading and unreliable. As shown in the Supporting Information (Section S7, Figures S9-S20), the RSV exhibits significantly high relative force estimation errors at low and high SNRs and reaches a minimum error in the vicinity of 0 dB where the variance contribution of downsampling and colored noises or drift cancel each other. This renders the RSV method unreliable as a force calibration method. For Brownian noise, linear and non-linear drift, the HV method results in the smallest mean normalized force estimation errors compared to the PSD and AV (Figures 4D,E). At high force (12 pN), the PSD, AV, and HV methods perform much better than the RSV method, which is explained by the dominant effects of downsampling on the accuracy of the RSV method, for which the PSD, AV, and HV are approximately invariant at sampling frequencies larger than the Nyquist frequency. Note that out of the three methods, the HV clearly performs best in presence of Brownian noise, linear, and non-linear drift, while performing similarly to PSD and AV for white and pink noise.

Together, our simulations show that the HV method is indeed the most stable, accurate, and precise in the presence of drift and performs very well for the other colored noise types, too. In addition, the relative errors using the PSD, AV, and HV methods to determine the variances of the motion trajectories decrease in the presence of white, pink and Brownian noise with increasing signal-to-noise ratios, which is expected for decreasing noise levels.

To further confirm the suitability of the HV as an accurate and drift-invariant method for force calibration in MT experiments, we apply the three methods to experimental data traces.

### Application of PSD, AV, and HV to experimental data traces

Measurements are carried out using a 21 kbp dsDNA construct and MyOne (diameter ≈1 μm) beads. As sampling frequencies, we choose a rate that fulfills the Nyquist theorem to minimize downsampling effects, i.e. 72 Hz. The position of the external magnet is gradually changed over a wide translation range. At each magnet position the bead motion is tracked for several tens of seconds (Figure 5A). We calculate the forces from the tracked bead positions for each magnet height using all three methods, PSD, AV, and HV (Figure 5B,C). A typical force calibration curve is shown in Figure 5D. The force decreases approximately exponentially with increasing magnet height, similar to the magnetic field^37^. Furthermore, the extension of the DNA tether is well-described by the 7-parameter WLC model (Figure 5E). From the WLC fit, we find a contour (*L*_*c*_) and persistence (*L*_*p*_) lengths of 6.8 μm and 44 nm, respectively, which are in good agreement with the expected crystallographic length of a ≈21 kbp DNA construct and with the literature values^37, 62, 72^ for *L*_*p*_ = 43 − 50 nm.

**Figure 5:**
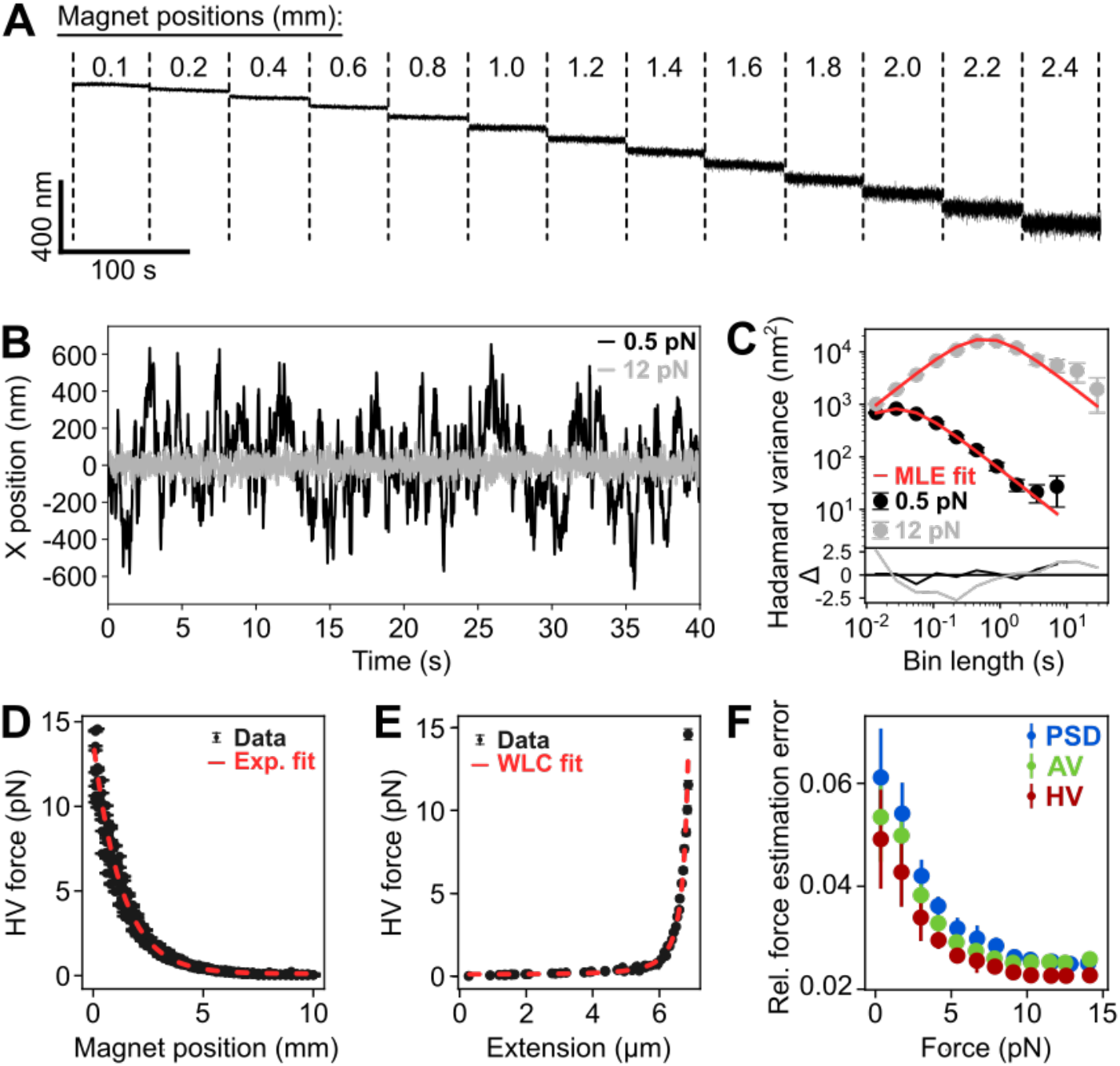
Force calibration using experimental MT data. **A)** Experimental extension time-traces (z-dimension) at different magnet positions corresponding to different forces. **B)** Experimental data traces (x-dimension) at low (0.5 pN) and high (12 pN) force. **C)** Hadamard variance of the experimental traces shown in B at low (0.5 pN) and high (12 pN) force. The red lines represent the MLE fits. **D)** HV force vs. magnet position plot of 13 beads. From the exponential fit (red dashed line), we find a maximum force of 13.9 pN. **E)** HV force-extension plot of one bead. The contour and persistence lengths derived from the WLC fit (red dashed line) are 6.8 μm and 44 nm, respectively. **F)** Relative force estimation errors of experimental data for PSD (blue), AV (green), and HV (red). Data are derived from experimental force calibration traces of 13 MyOne beads (diameter: ≈1 μm, sampling frequency: 72 Hz).

Next, we determine the standard force estimation error from the covariance of the maximum likelihood estimation (MLE) fit to the experimental data (Supporting Information, Section S8). Note that we do not know the ground truth of the forces that we calculate from experimental traces, making it difficult to confirm which method performs best in practice. However, one can examine how well each method is internally consistent, i.e how well the MLE fits match the experimental data, which we use as a metric for confirming the accuracy of a method. Therefore, we calculate the relative force estimation errors (standard force estimation error over force) as a function of the force. We bin the relative errors to 12 force steps in total (Figure 5F). We find that over the whole force range the HV method results in lower relative errors for the studied bead-tether system than the PSD and AV. On average, the HV errors are 18.0 % and 9.4 % lower compared to the PSD and AV, respectively. A similar trend is found for the M270 beads (diameter ≈2.8 μm; Supporting information, Section S9, Figure S21). This finding indicates that the HV method is the most precise in analyzing the experimental data. As a control, we also analyze the relative bead-to-bead force errors and find a force uncertainty of ≈11 % which is in good agreement with previous reports^37, 44^ that report a bead-to-bead variability of approximately 10%, which is dominated by differences in bead volumes and magnetic particle content (Supporting Information, Section S10, Figure S22). Notably, this error is the same for all three methods demonstrating that the bead-to-bead variations are a systematic error of the system and independent of the applied method.

## Conclusion

In conclusion, we introduce a robust thermal motion-based method using the Hadamard variance, for processing and analyzing multiplexed magnetic tweezer data with high accuracy. By addressing common sources of additive noise in comparison to conventional methods such as the real-space variance, the power spectral density, and the Allan variance, we find that the HV method offers important improvements in force calibration precision. Through extensive testing with Brownian dynamics simulations, replicating typical experimental MT conditions, and including controlled levels of common colored noise types and drift, we demonstrate the superior performance of the HV method across a wide range of signal-to-noise ratios and drift speeds. Notably, our results show consistently lower force estimation errors with the HV method, particularly at SNR ≥ 10 dB, and high stability in the presence of drift, ensuring reliable results across varying experimental conditions. Application of the HV method to experimental datasets further validates its effectiveness, revealing an average reduction of force estimation errors by 18.0 % and 9.4 % compared to the PSD and AV methods, respectively. These findings highlight the suitability of the HV method for accurate force calibration, which is important for achieving sub-pN resolution and precision in multiplexed MT experiments. In summary, our study contributes a new force calibration method to the existing repertoire, offering researchers a powerful tool for obtaining accurate results that are essential for advancing our understanding of molecular-scale interactions and mechanical processes in biological systems. For this purpose, we make our approach widely accessible *via* a clearly documented open-source implementation of the HV method as an extension of the existing *Tweezepy*^67^ package.

## ACKNOWLEDGEMENTS

This work was supported by the European Research Council through the Consolidator Grant “ProForce” and the Dutch Research Council (NWO) with grant number OCENW.GROOT.2019.071. SDP was further supported by the Alexander-von-Humboldt foundation through a Feodor-Lynen fellowship. The authors thank Dave van den Heuvel and Elleke van Harten for laboratory assistance.

