## Supplementary Information for "Accurate Drift-Invariant Single-Molecule Force Calibration Using the Hadamard Variance"

June 2024

### Contents

**S1 Closed Form Expression of Hadamard Variance of a Magnetic Bead in a Magnetic Trap 2**

**S2 Colored Noise Contributions in Experimental MT Data Traces. 3**

**S3 Relative Force Estimation Errors in Presence of Downsampling 5**

**S4 Downsampling Correction Efficiency of the Power Spectral Density, Allan Variance, Hadamard Variance, and Real-Space Variance 6**

**S5 Relative Force Estimation Errors in Presence of Colored Noise and Non-linear Drift 7**

**S6 Relative Force Estimation Errors in Presence of Linear Drift as a Function of Drift Speed. 11**

**S7 Application of the Real-Space Variance Method to Simulation Traces in Presence of Downsampling, Colored Noise, and Linear and Non-Linear Drift 12**

**S8 Maximum Likelihood Estimation Fitting 21**

**S9 Experimental Force Estimation Errors of M270 beads 22**

**S10 Experimental Relative Bead-to-Bead Force Errors as a Function of the Force for MyOne Beads 22**

**References 23**

### S1 Closed Form Expression of Hadamard Variance of a Magnetic Bead in a Magnetic Trap

In this section, we derive the closed form expression of the Hadamard variance of a magnetic bead in a magnetic trap. The motion of a tethered paramagnetic bead in a magnetic tweezer setup around the equilibrium is well described by the Ornstein-Uhlenbeck (OU) process, i.e., Brownian motion within a harmonic trap. Without loss of generality, we focus on the motion in the $x$-axis, and the equation of motion is given by[3]:

$$\gamma \frac{\mathrm{d}x(t)}{\mathrm{d}t} + \kappa x(t) = F_L(t), \tag{1}$$

where $x(t)$ denotes the bead position in $x$-axis, $\kappa$ is the stiffness of the harmonic trap formed by the tether-magnet system, $\gamma$ is the Stoke's drag coefficient, and $F_L(t)$ is the stochastic Langevin force which follows the fluctuation-dissipation relation $\langle F_L(t)F_L(t')\rangle = 2\gamma k_B T\delta(t-t')$.

The OU process is well-studied and its power spectral density admits the closed-form[3]

$$S(f) = \frac{k_B T}{2\pi^2\gamma f_c^2}\frac{1}{1+(f/f_c)^2}, \tag{2}$$

where $k_B$ is the Boltzmann constant, $T$ is the temperature, and the corner frequency is defined by $f_c \equiv \kappa/2\pi\gamma$. Having a closed-form expression for the power spectral density plays a crucial role for our Hadamard variance (HV) based force calibration technique since we can express HV in terms of it as [8]

$$\sigma_H^2(\tau) = \int_0^\infty |H_H(f,\tau)|^2 S(f)\mathrm{d}f, \tag{3}$$

where the transfer function $|H_H(f,\tau)|^2$ is given by [1]

$$|H_H(f,\tau)|^2 = \frac{16}{3}\frac{\sin^6(\pi f\tau)}{(\pi f\tau)^2}. \tag{4}$$

Here, we present a brief derivation of the closed form of HV for our force calibration procedure. We start by pluging Eq.(2) and Eq.(4) into Eq.(3) and use the symmetry around the $f=0$ point of the integrant to symmetrize the integral. This yields

$$\sigma_H^2(\tau) = \frac{4k_B T}{3\tau\pi^3\gamma f_c^2}\int_{-\infty}^\infty \frac{\sin^6(\xi)}{\xi^2\left[1+(\xi/\ell)^2\right]}\mathrm{d}\xi, \tag{5}$$

where we introduce the dimensionless variables $\xi \equiv \pi f\tau$ and $\ell \equiv \pi\tau f_c$. To put the integral into a familiar form, we introduce the Binomial expansion of the $\sin^6(\xi)$ term,

$$\sin^6(\xi) = \left(\frac{e^{i\xi}-e^{-i\xi}}{2i}\right)^6 = \frac{1}{64}\sum_{n=-3}^{3}\binom{6}{n+3}(-1)^n e^{-i2n\xi}. \tag{6}$$

Using this identity, HV takes the form

$$\sigma_H^2(\tau) = \frac{k_B T}{48\tau\pi^3\gamma f_c^2}\sum_{n=-3}^{3}\binom{6}{n+3}(-1)^n\int_{-\infty}^\infty \frac{e^{-i2n\xi}}{\xi^2\left[1+(\xi/\ell)^2\right]}\mathrm{d}\xi. \tag{7}$$

Now, notice that with defining the function $s(\xi) \equiv 1/\xi^2[1+(\xi/\ell)^2]$, the integral term in HV is just a Fourier transform,

$$\tilde{s}(\omega) = \int_{-\infty}^\infty s(\xi)e^{-i\omega\xi}\mathrm{d}\xi \tag{8}$$

$$= \int_{-\infty}^\infty \frac{e^{-i\omega\xi}}{\xi^2\left[1+(\xi/\ell)^2\right]}\mathrm{d}\xi \tag{9}$$

$$= -\pi|\omega| - \frac{\pi}{\ell}e^{-\ell|\omega|}. \tag{10}$$

Therefore, HV becomes

$$\sigma_H^2(\tau) = \frac{k_B T}{48\tau\pi^3\gamma f_c^2}\sum_{n=-3}^{3}\binom{6}{n+3}(-1)^n\tilde{s}(2n), \tag{11}$$

and evaluating the sum and plugging back the original variables yields the final expression for HV,

$$\sigma_H^2(\tau) = \frac{2\gamma k_B T}{\kappa^2 \tau}\left(1 - \frac{5\gamma}{3\kappa\tau} + \frac{5\gamma}{2\kappa\tau}e^{-\frac{\kappa\tau}{\gamma}} - \frac{\gamma}{\kappa\tau}e^{-\frac{2\kappa\tau}{\gamma}} + \frac{\gamma}{6\kappa\tau}e^{-\frac{3\kappa\tau}{\gamma}}\right). \tag{12}$$

Just like Allan variance, Hadamard variance intrinsically accounts for downsampling errors since its calculation from the experimental measuerements is based on averaged positions. On the other hand, sampling frequency ($f_s$) of the magnetic tweezer setup sets the lower bound of the observation time $\tau$ such that $\tau \geq 1/f_s$.

### S2 Colored Noise Contributions in Experimental MT Data Traces.

In order to identify the most common colored noise types in our experimental MT data, we determine the power spectral density (PSD), Allan variance (AV), and HV of immobilized reference beads as follows: We identify reference bead traces that have a common trend of the overall mechanical drift. The identification is done as follows:

1. We calculate the first forward difference of the reference bead candidates. This ensures the identification process stays translationally invariant.
2. We apply exponential smoothing [2, 4, 10] on the first forward differences to suppress the noise contamination. This gives us the smooth trend of the reference bead candidates.
3. We apply the isolation forest algorithm [5, 6] to eliminate outlying candidates.
4. The remaining reference bead candidates form the set of reference beads.

Next, we calculate the overall trend of the reference bead. The procedure we use is the following:

1. We shift every reference bead trace by its mean value to center the trace around zero.
2. We calculate the average trace of the set of reference beads.
3. We apply exponential smoothing to the average trace to suppress the remaining noise.
4. We end up with the overall smooth trend of the reference beads.

After obtaining the overall trend, we detrend the reference beads by subtracting the overall trend from every reference bead. Since reference beads are immobilized, the resulting detrended traces contain only noise. Therefore, we estimate the contribution of different types of noise by calculating the HV of each detrended reference trace and fit the data for the contributions of white noise ($h_0$), pink noise ($h_{-1}$), and Brownian noise ($h_{-2}$) using [8]

$$\sigma^2_{\text{HV,noise}} = \underbrace{\frac{h_0}{2\tau}}_{\text{White noise}} + \underbrace{\frac{1}{2}\log\left(\frac{256}{27}\right)h_{-1}}_{\text{Pink noise}} + \underbrace{\frac{\pi^2\tau}{3}h_{-2}}_{\text{Brown noise}} \tag{13}$$

with $\tau$ being the bin length.

For completeness, we also calculate the PSD and AV of the detrended reference traces and plot the corresponding noise curves using the contribution parameters that we obtained from HV. The noise curves of AV and PSD are given by [8]

$$\sigma^2_{\text{AV,noise}} = \underbrace{\frac{h_0}{2\tau}}_{\text{White noise}} + \underbrace{2\log(2)\,h_{-1}}_{\text{Pink noise}} + \underbrace{\frac{2\pi^2\tau}{3}h_{-2}}_{\text{Brown noise}}, \tag{14}$$

and

$$S(f) = \underbrace{h_0}_{\text{White noise}} + \underbrace{\frac{h_{-1}}{f}}_{\text{Pink noise}} + \underbrace{\frac{h_{-2}}{f^2}}_{\text{Brown noise}}. \tag{15}$$

with $f$ being the frequency. As shown in Fig. S1, the PSD, and AV/HV variances are well described by the total noise curve, i.e. the sum of the white, pink, and Brownian noise contributions.

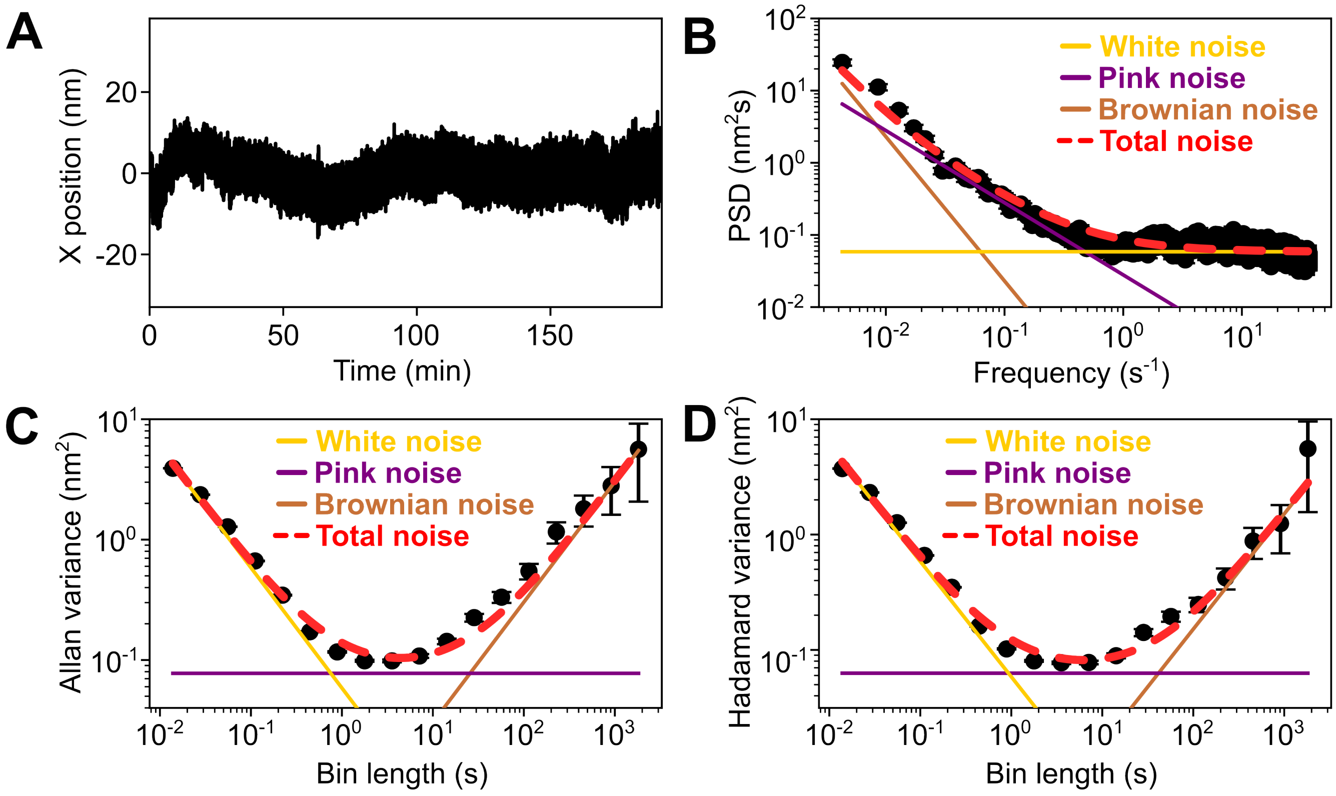


**Figure S1:** Contributions of colored noise in experimental MT traces. **(A)** Experimental reference bead trace after drift correction. **(B)** PSD of the experimental reference bead trace as a function of the frequency. **(C)** AV of the experimental reference bead trace as a function of the bin length. **(D)** HV of the experimental reference bead trace as a function of the bin length. The yellow, pink, and brown solid lines represent white, pink, and Brownian noise, respectively. The dotted red lines represent the total noise, i.e. the sum of the white, pink, and Brownian noise contributions.

### S3 Relative Force Estimation Errors in Presence of Downsampling

We conduct Brownian dynamics simulations of 1 $\mu$m beads, similar to MyOne beads, at a force of 2.5 pN, as detailed in the main text. Additionally, we simulate 3 $\mu$m beads, similar to M270 beads, at forces of 0.5, 2.5, 12, 40, and 80 pN. We analyze the relative force estimation errors at different sampling frequencies using the PSD, AV, and HV methods. As illustrated in Fig. S2, the relative force estimation errors increase as the sampling frequency decreases in all cases, attributable to stronger downsampling.

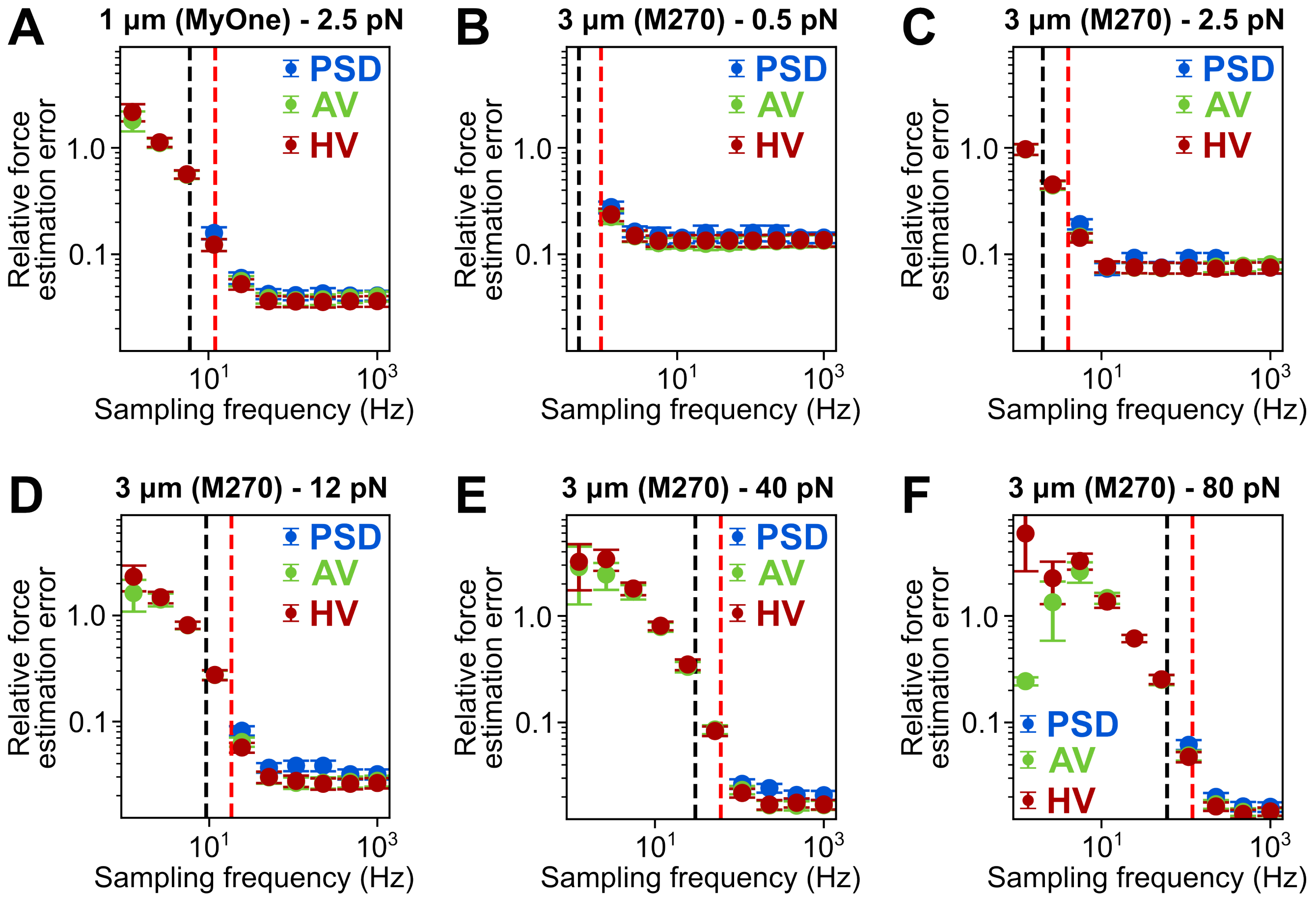


**Figure S2:** Effect of downsampling on the force estimation using simulated data traces. **(A)** Relative force estimation errors as a function of the sampling frequency for PSD (blue), AV (green), and HV (red) of 1 $\mu$m beads resembling MyOne magnetic beads at a force of 2.5 pN. **(B-F)** Relative force estimation errors as a function of the sampling frequency for PSD (blue), AV (green), and HV (red) of 3 $\mu$m beads (mimicking M270) at forces of 0.5 pN (B), 2.5 pN (C), 12 pN (D), 40 pN (E), and 80 pN (F). The black and red dashed lines represent the corner and Nyquist frequencies, respectively.

#### S4 Downsampling Correction Efficiency of the Power Spectral Density, Allan Variance, Hadamard Variance, and Real-Space Variance

In order to quantify the downsampling correction efficiency of the PSD, AV, HV, and real-space variance (RSV) methods, we simulate the Brownian dynamics of 1 $\mu$m beads, similar to MyOne beads, at forces ranging from 0.5 pN to 12 pN (4 simulations per force), as described in the main text. After downsampling the simulated traces to match 72 Hz sampling rate, we use PSD, AV, HV, and RSV methods to estimate the force acting on the bead. Finally, we plot the estimated force values as a function of the *true* force that we used to simulate the trajectories. We observe that all spectral methods, i.e. PSD, AV, and HV, perform with similar accuracy while RSV performs significantly worse.

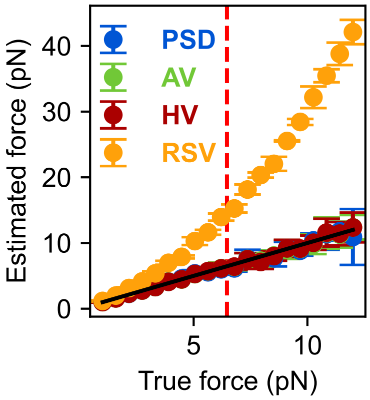


**Figure S3:** Downsampling correction efficiency for the PSD (blue), AV(green), HV (red) and RSV (orange) methods at forces ranging from 0.5 pN to 12 pN. The sampling frequency is set to 72 Hz and the red dashed line represents the force that corresponds to the Nyquist frequency $f_N = 2f_c = 72$ Hz. The black solid line represents the ideal relation between estimated and *true* force, i.e. estimated force = *true* force.

#### S5 Relative Force Estimation Errors in Presence of Colored Noise and Non-linear Drift

We perform Brownian dynamics simulations using 1 $\mu$m beads (mimicking MyOne) at a force of 2.5 pN and 3 $\mu$m beads (mimicking M270) at forces of 0.5, 2.5, 12, 40, and 80 pN, and add white noise (Fig. S4), pink noise (Fig. S5), and Brownian noise (Fig. S6) as well as non-linear drift (Fig. S7), as detailed in the main text. We analyze the relative force estimation errors as a function of the signal-to-noise ratios (SNR) using the PSD, AV, and HV methods. The sampling frequency is 72 Hz.

##### White noise

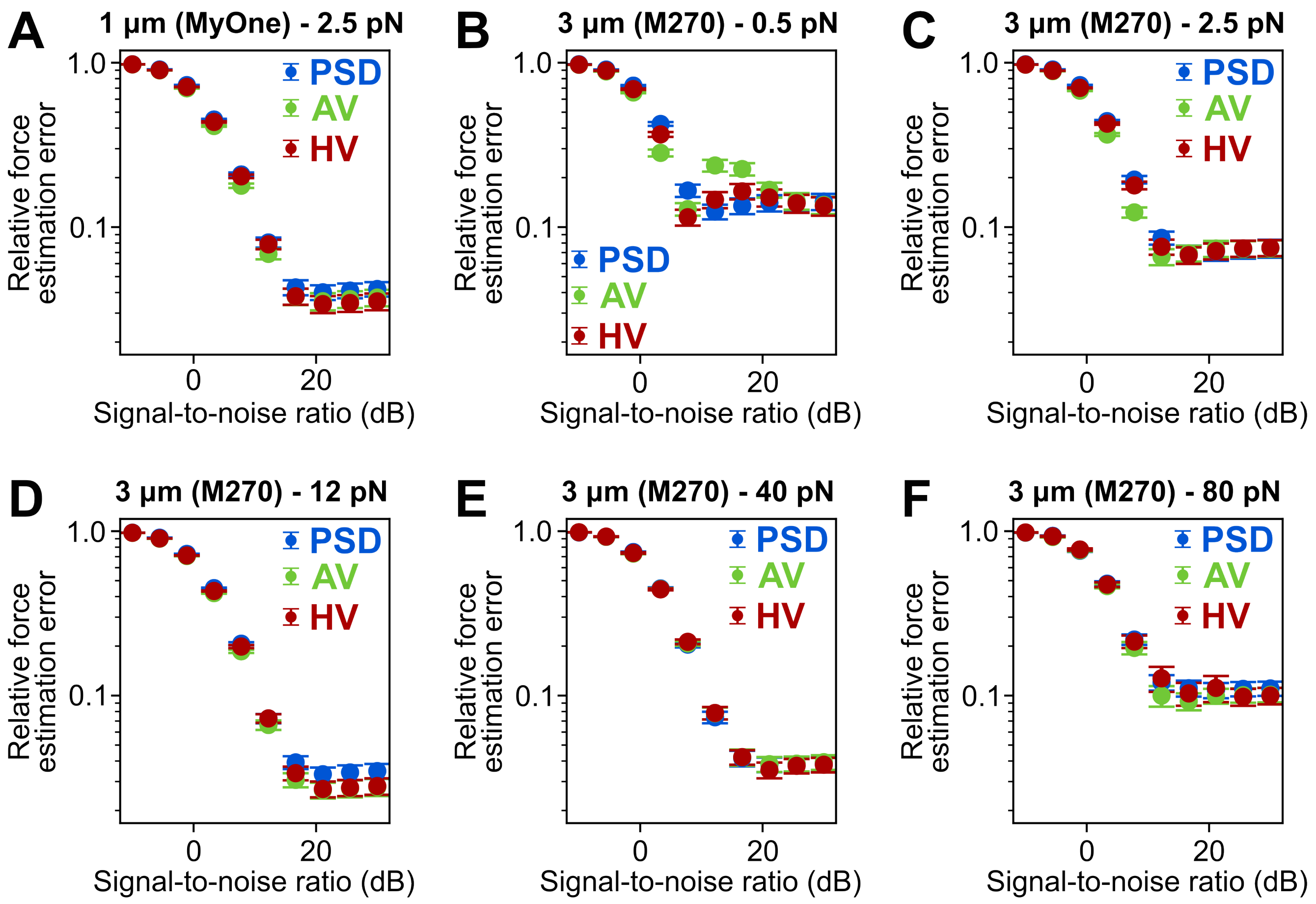


**Figure S4:** Effect of white noise on the force estimation errors. **(A)** Relative force estimation errors as a function of the signal-to-noise ratio for PSD (blue), AV (green), and HV (red) of 1 $\mu$m beads at a force of 2.5 pN. **(B-F)** Relative force estimation errors as a function of the signal-to-noise ratio for PSD (blue), AV (green), and HV (red) of 3 $\mu$m beads at forces of 0.5 pN (B), 2.5 pN (C), 12 pN (D), 40 pN (E), and 80 pN (F).

#### Pink noise

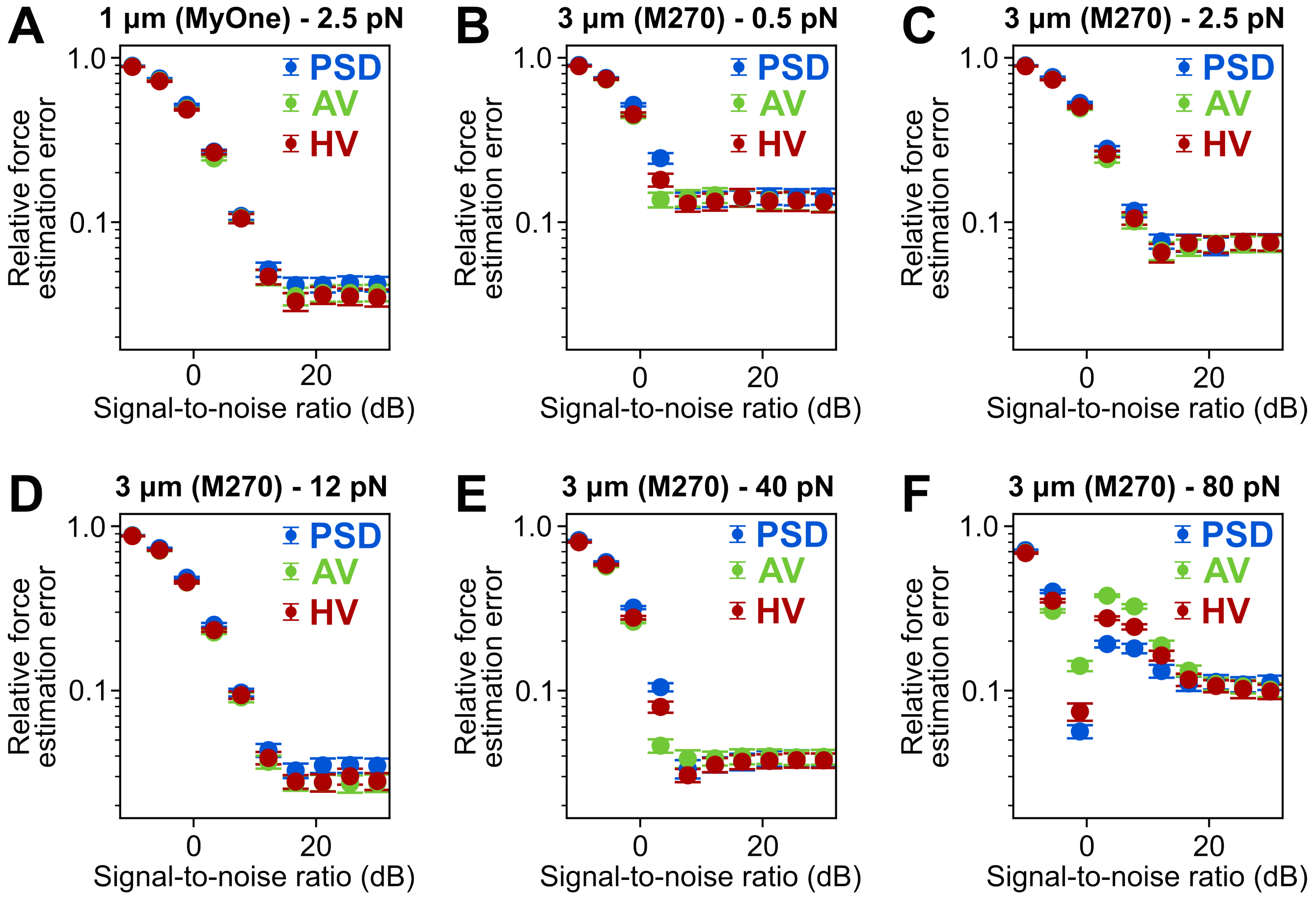


**Figure S5:** Effect of pink noise on the force estimation errors. **(A)** Relative force estimation errors as a function of the signal-to-noise ratio for PSD (blue), AV (green), and HV (red) of 1 $\mu$m beads at a force of 2.5 pN. **(B-F)** Relative force estimation errors as a function of the signal-to-noise ratio for PSD (blue), AV (green), and HV (red) of 3 $\mu$m beads at forces of 0.5 pN (B), 2.5 pN (C), 12 pN (D), 40 pN (E), and 80 pN (F).

<u>Brownian noise</u>

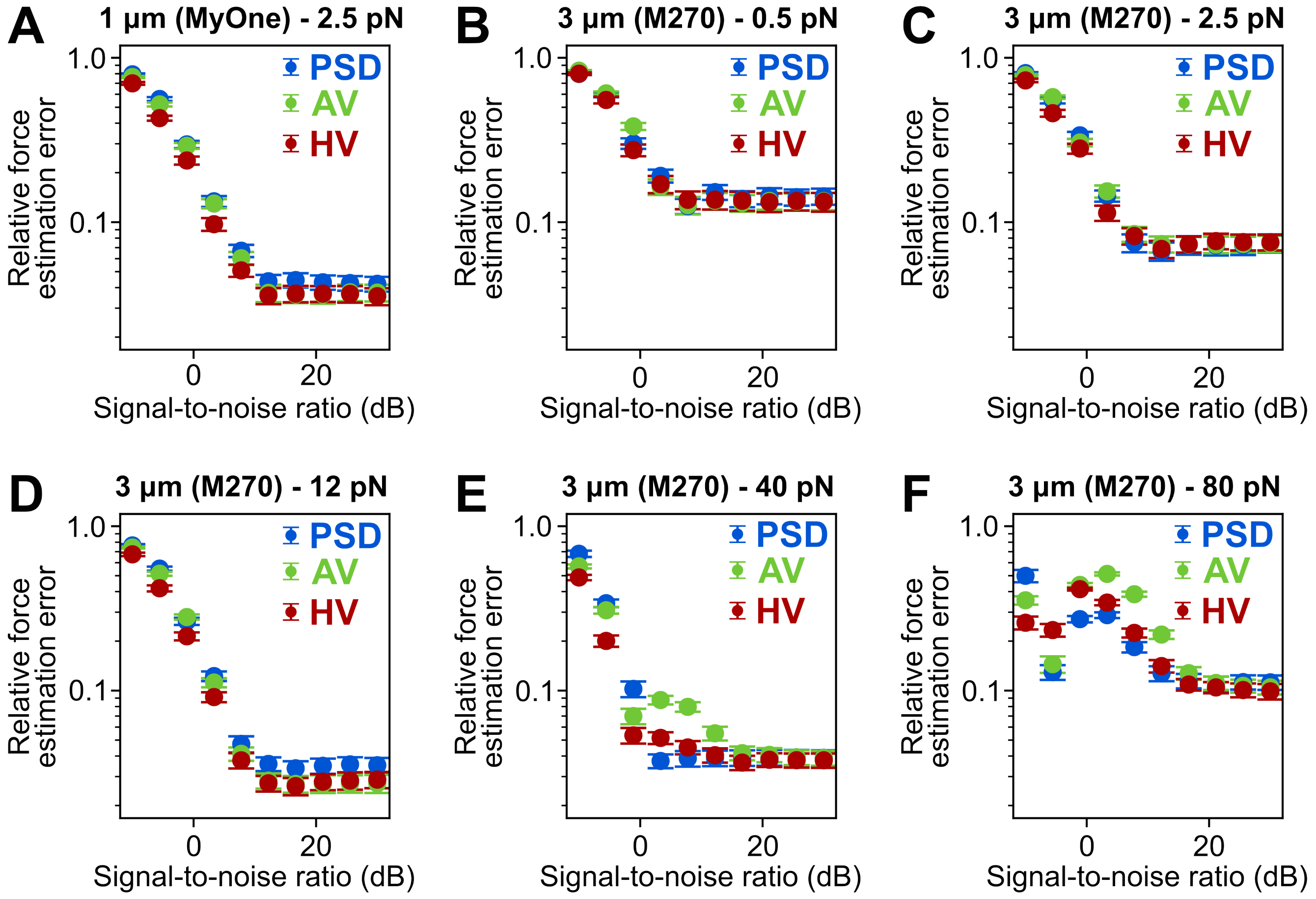


**Figure S6:** Effect of Brownian noise on the force estimation errors. **(A)** Relative force estimation errors as a function of the signal-to-noise ratio for PSD (blue), AV (green), and HV (red) of 1 $\mu$m beads at a force of 2.5 pN. **(B-F)** Relative force estimation errors as a function of the signal-to-noise ratio for PSD (blue), AV (green), and HV (red) of 3 $\mu$m beads at forces of 0.5 pN (B), 2.5 pN (C), 12 pN (D), 40 pN (E), and 80 pN (F).

### Non-linear drift

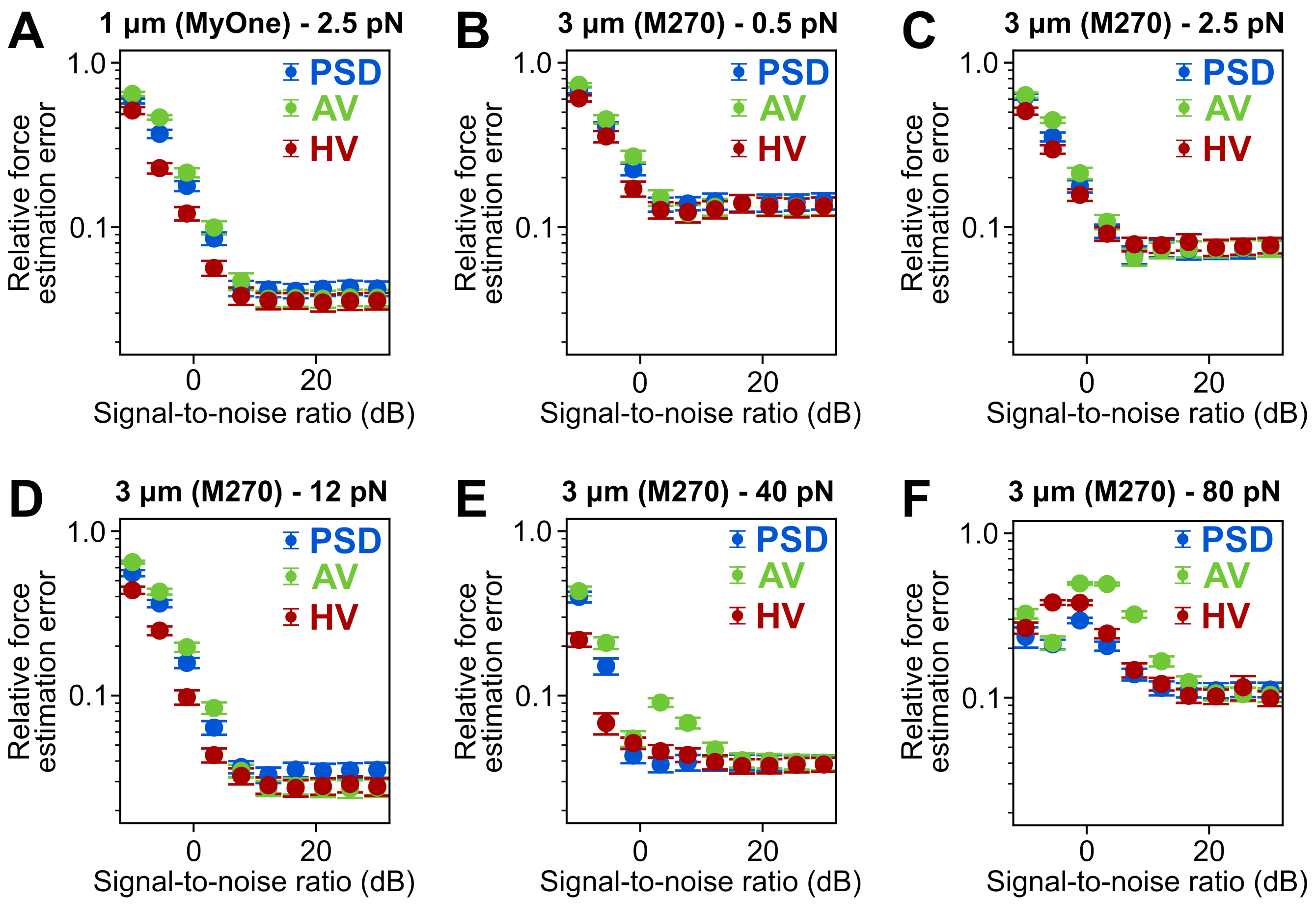


**Figure S7:** Effect of non-linear drift on the force estimation errors. **(A)** Relative force estimation errors as a function of the signal-to-noise ratio for PSD (blue), AV (green), and HV (red) of 1 $\mu$m beads at a force of 2.5 pN. **(B-F)** Relative force estimation errors as a function of the signal-to-noise ratio for PSD (blue), AV (green), and HV (red) of 3 $\mu$m beads at forces of 0.5 pN (B), 2.5 pN (C), 12 pN (D), 40 pN (E), and 80 pN (F).

### S6 Relative Force Estimation Errors in Presence of Linear Drift as a Function of Drift Speed.

We perform Brownian dynamics simulations using 1 $\mu$m beads (mimicking MyOne) at a force of 2.5 pN and 3 $\mu$m beads (mimicking M270) at forces of 0.5, 2.5, 12, 40, and 80 pN, and add linear drift to the traces as detailed in the main text. We analyze the relative force estimation errors as a function of the drift speed using the PSD, AV, and HV methods. The sampling frequency is 72 Hz.

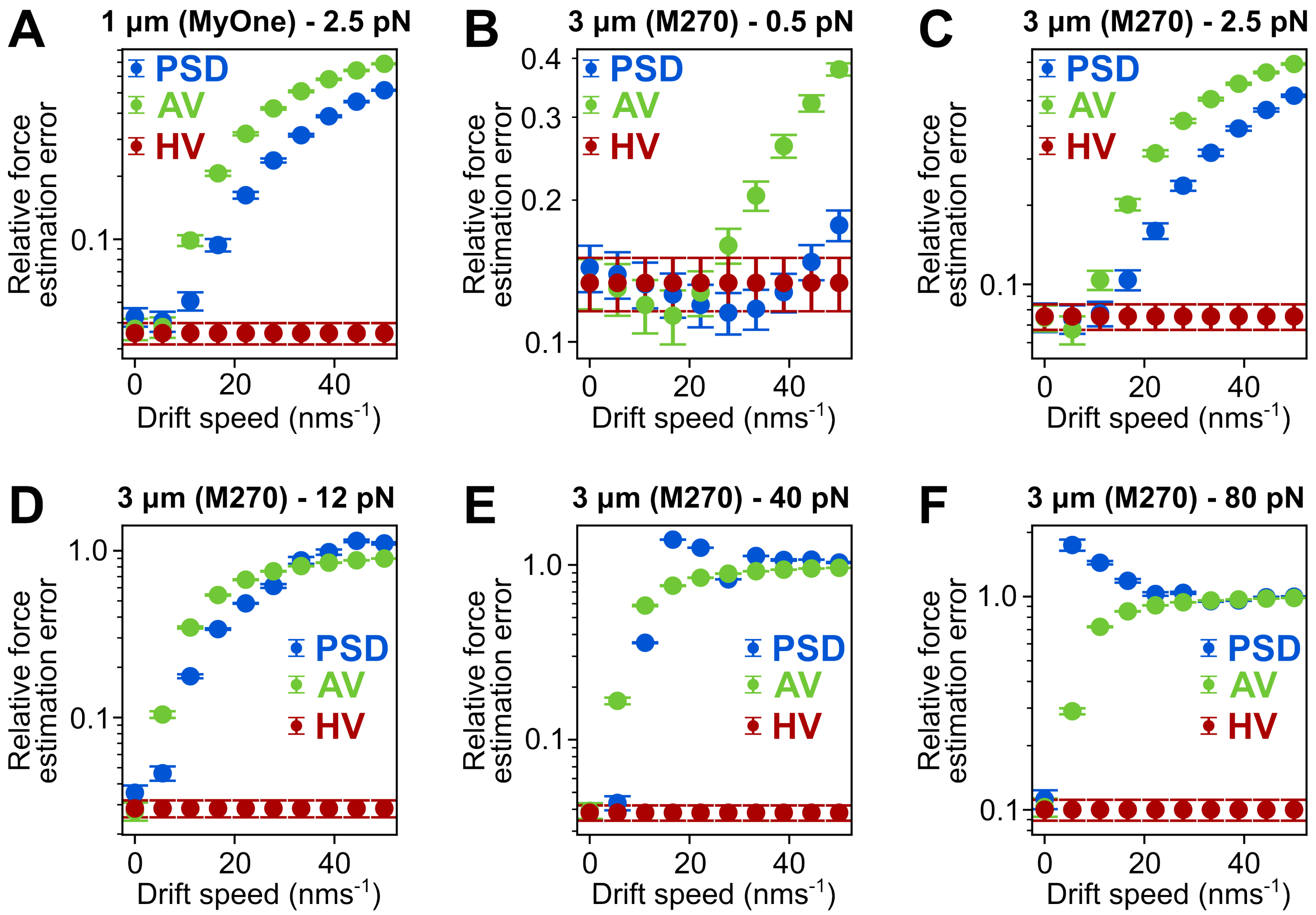


**Figure S8:** Effect of linear drift on the force estimation errors. **A)** Relative force estimation errors as a function of the drift speed for PSD (blue), AV (green), and HV (red) of 1 $\mu$m beads at a force of 2.5 pN. **B-F)** Relative force estimation errors as a function of the drift speed for PSD (blue), AV (green), and HV (red) of 3 $\mu$m beads at forces of 0.5 pN (B), 2.5 pN (C), 12 pN (D), 40 pN (E), and 80 pN (F).

### S7 Application of the Real-Space Variance Method to Simulation Traces in Presence of Downsampling, Colored Noise, and Linear and Non-Linear Drift

In order to study the real-space variance (RSV) method in presence of downsampling, colored noise and linear and non-linear drift, we perform Brownian dynamics simulations using 1 $\mu$m beads (mimicking MyOne) at forces of 0.5, 2.5, and 12 pN, and 3 $\mu$m beads (mimicking M270) at forces of 0.5, 2.5, 12, 40, and 80 pN, as detailed in the main text. The DNA-tether is again modeled as a worm-like-chain whose contour and persistence length are 7 $\mu$m (corresponding to 21 kbp dsDNA) and 45 nm, respectively. The effects of downsampling, white, pink, Brownian noise, and linear and non-linear drift for the 1 $\mu$m and 3 $\mu$m beads are shown in Figs. S9-S20, respectivley.

**1 $\mu$m (MyOne) beads - downsampling**

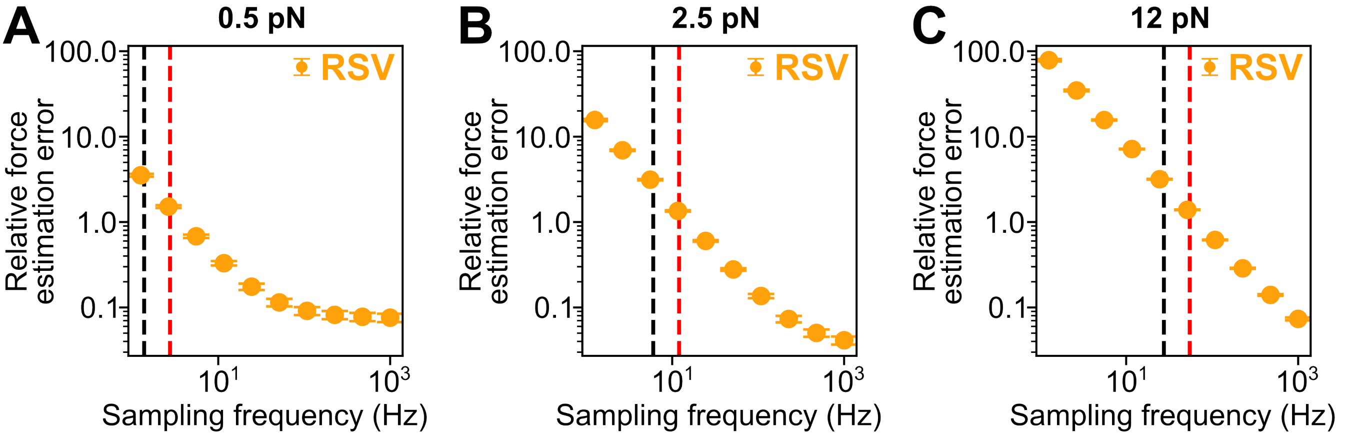


**Figure S9:** Effect of downsampling on the force estimation errors evaluated using simulated traces of 1 $\mu$m beads. Relative force estimation errors as a function of the sampling frequency for RSV at **(A)** 0.5 pN, **(B)** 2.5 pN, and **(C)** 12 pN. The black dashed lines highlight the corner frequencies and the red dashed lines the Nyquist frequencies.

**1 $\mu$m (MyOne) beads - white noise**

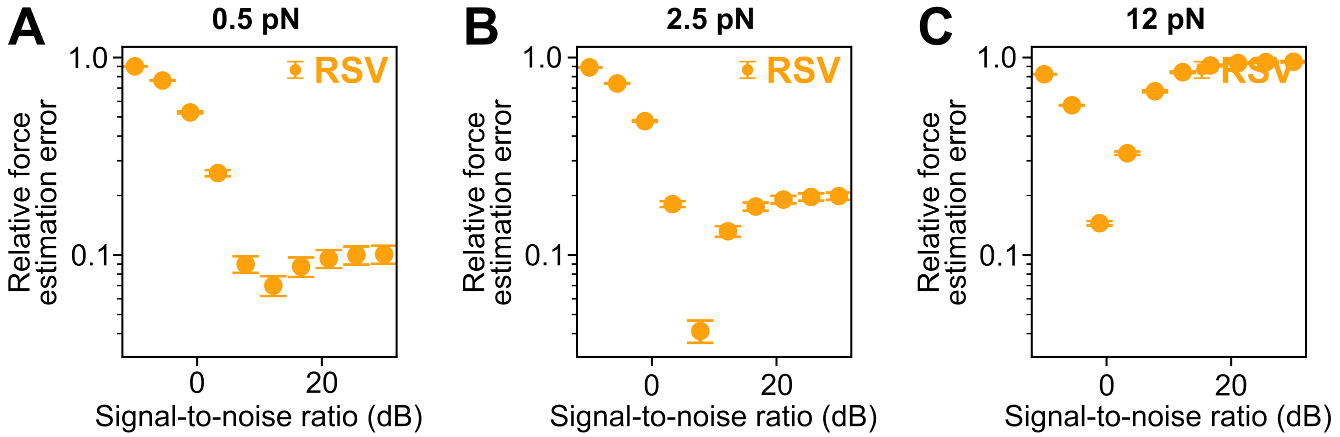


**Figure S10:** Effect of white noise on the force estimation errors evaluated using simulated traces of 1 $\mu$m beads. Relative force estimation errors as a function of the signal-to-noise ratio for RSV at **(A)** 0.5 pN, **(B)** 2.5 pN, and **(C)** 12 pN. The sampling frequency is 72 Hz.

#### $1\,\mu$m (MyOne) beads - pink noise

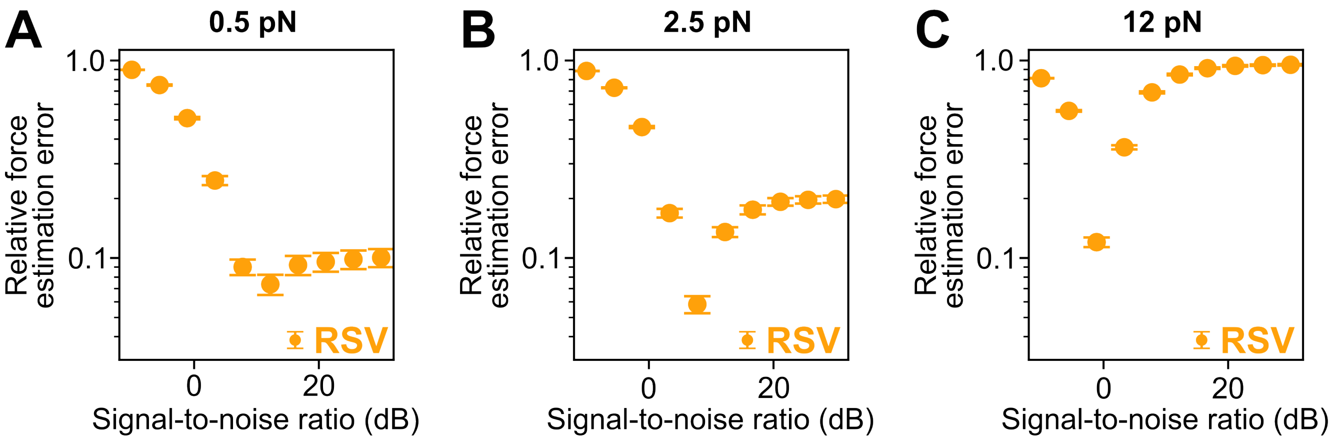


**Figure S11:** Effect of pink noise on the force estimation errors evaluated using simulated traces of $1\,\mu$m beads. Relative force estimation errors as a function of the signal-to-noise ratio for RSV at **(A)** 0.5 pN, **(B)** 2.5 pN, and **(C)** 12 pN. The sampling frequency is 72 Hz.

#### $1\,\mu$m (MyOne) beads - Brownian noise

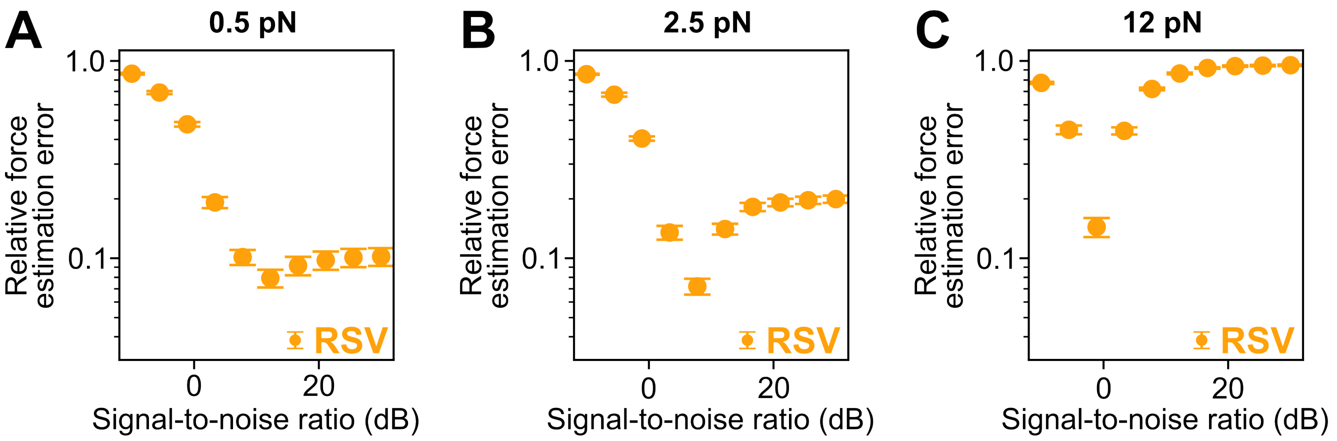


**Figure S12:** Effect of Brownian noise on the force estimation errors evaluated using simulated traces of $1\,\mu$m beads. Relative force estimation errors as a function of the signal-to-noise ratio for RSV at **(A)** 0.5 pN, **(B)** 2.5 pN, and **(C)** 12 pN. The sampling frequency is 72 Hz.

#### $1\,\mu$m (MyOne) beads - linear drift

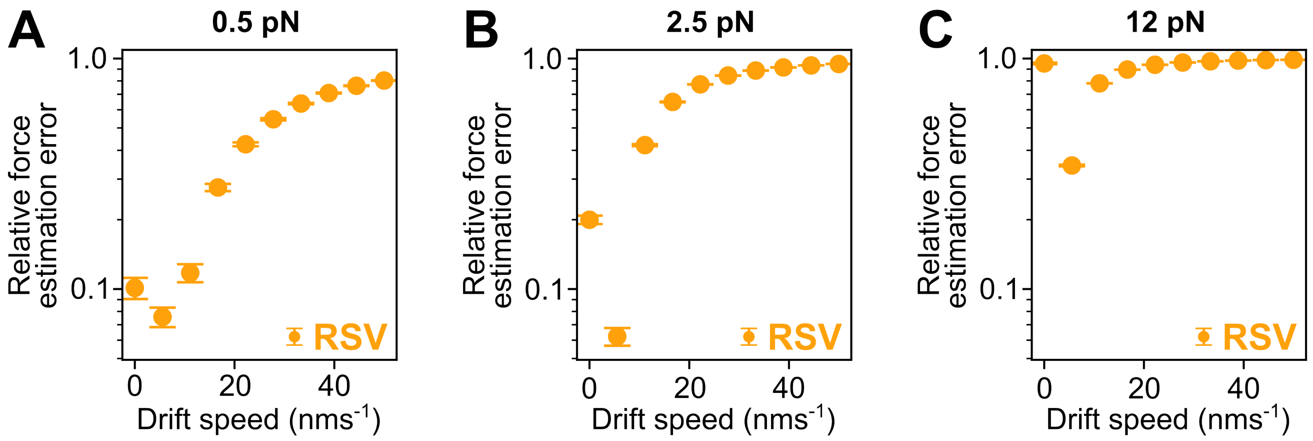


**Figure S13:** Effect of linear drift on the force estimation errors evaluated using simulated traces of $1\,\mu$m beads. Relative force estimation errors as a function of the drift speed for RSV at **(A)** 0.5 pN, **(B)** 2.5 pN, and **(C)** 12 pN. The sampling frequency is 72 Hz.

#### $1\,\mu$m (MyOne) beads - non-linear drift

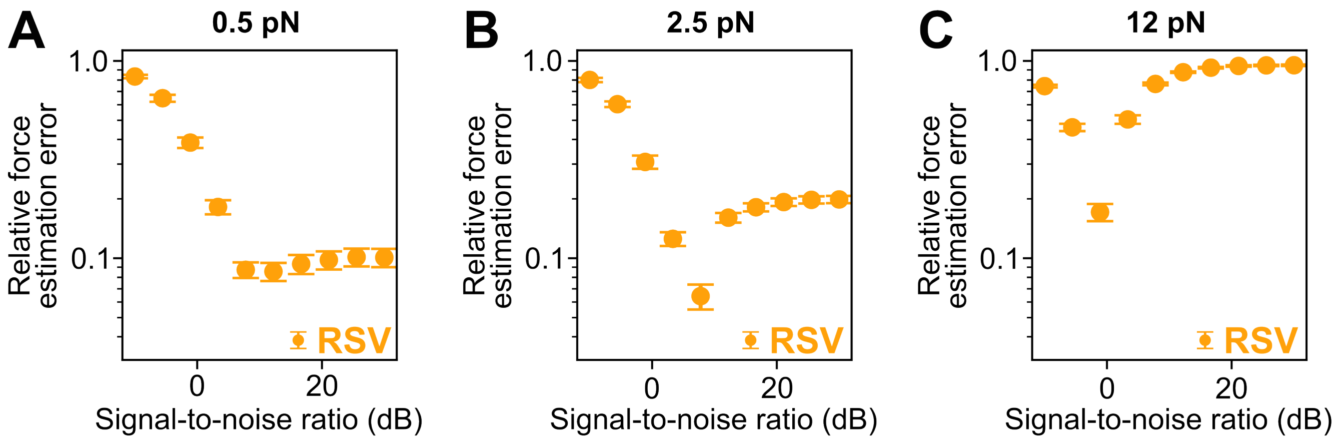


**Figure S14:** Effect of non-linear drift on the force estimation errors evaluated using simulated traces of $1\,\mu$m beads. Relative force estimation errors as a function of the signal-to-noise ratio for RSV at **(A)** 0.5 pN, **(B)** 2.5 pN, and **(C)** 12 pN. The sampling frequency is 72 Hz.

### 3 μm (M270) beads - downsampling

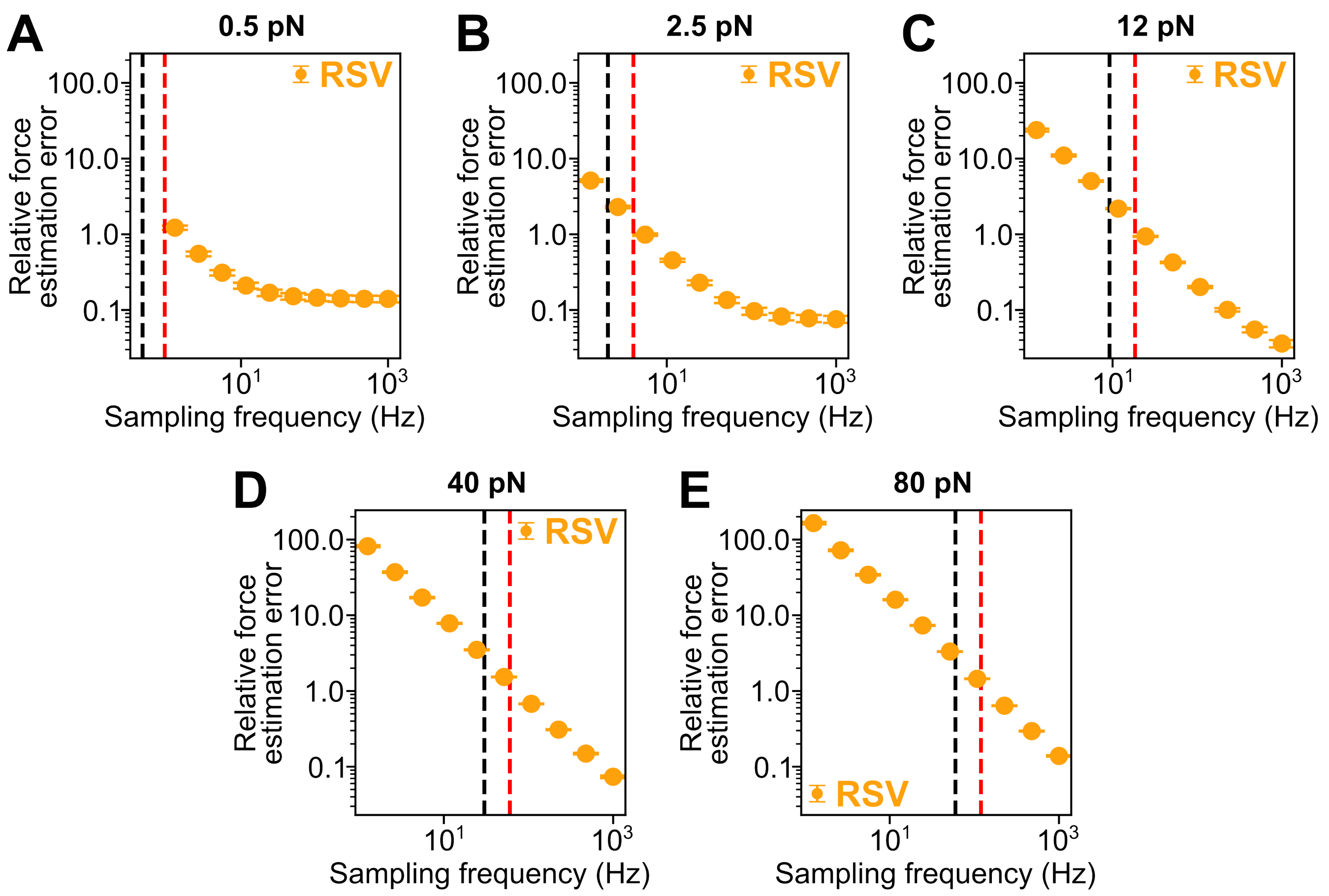


**Figure S15:** Effect of downsampling on the force estimation errors evaluated using simulated traces of 3 μm beads. Relative force estimation errors as a function of the sampling frequency for RSV at **(A)** 0.5 pN, **(B)** 2.5 pN, and **(C)** 12 pN, **(D)** 40 pN, and **(E)** 80 pN. The black dashed lines highlight the corner frequencies and the red dashed lines the Nyquist frequencies.

#### 3 $\mu$m (M270) beads - white noise

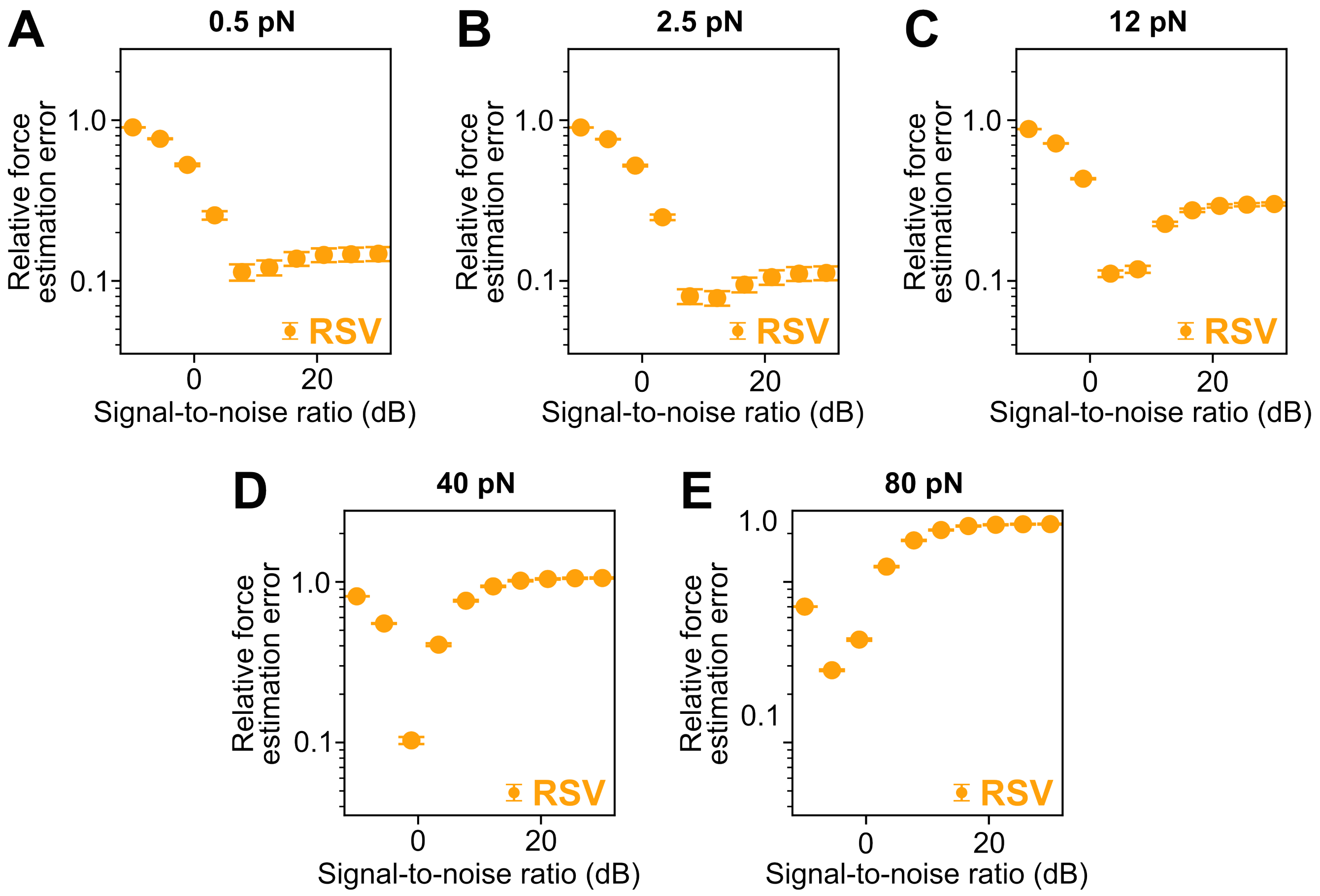


**Figure S16:** Effect of white noise on the force estimation errors evaluated using simulated traces of 3 $\mu$m beads. Relative force estimation errors as a function of the signal-to-noise ratio for RSV at **(A)** 0.5 pN, **(B)** 2.5 pN, and **(C)** 12 pN, **(D)** 40 pN, and **(E)** 80 pN. The sampling frequency is 72 Hz.

#### 3 $\mu$m (M270) beads - pink noise

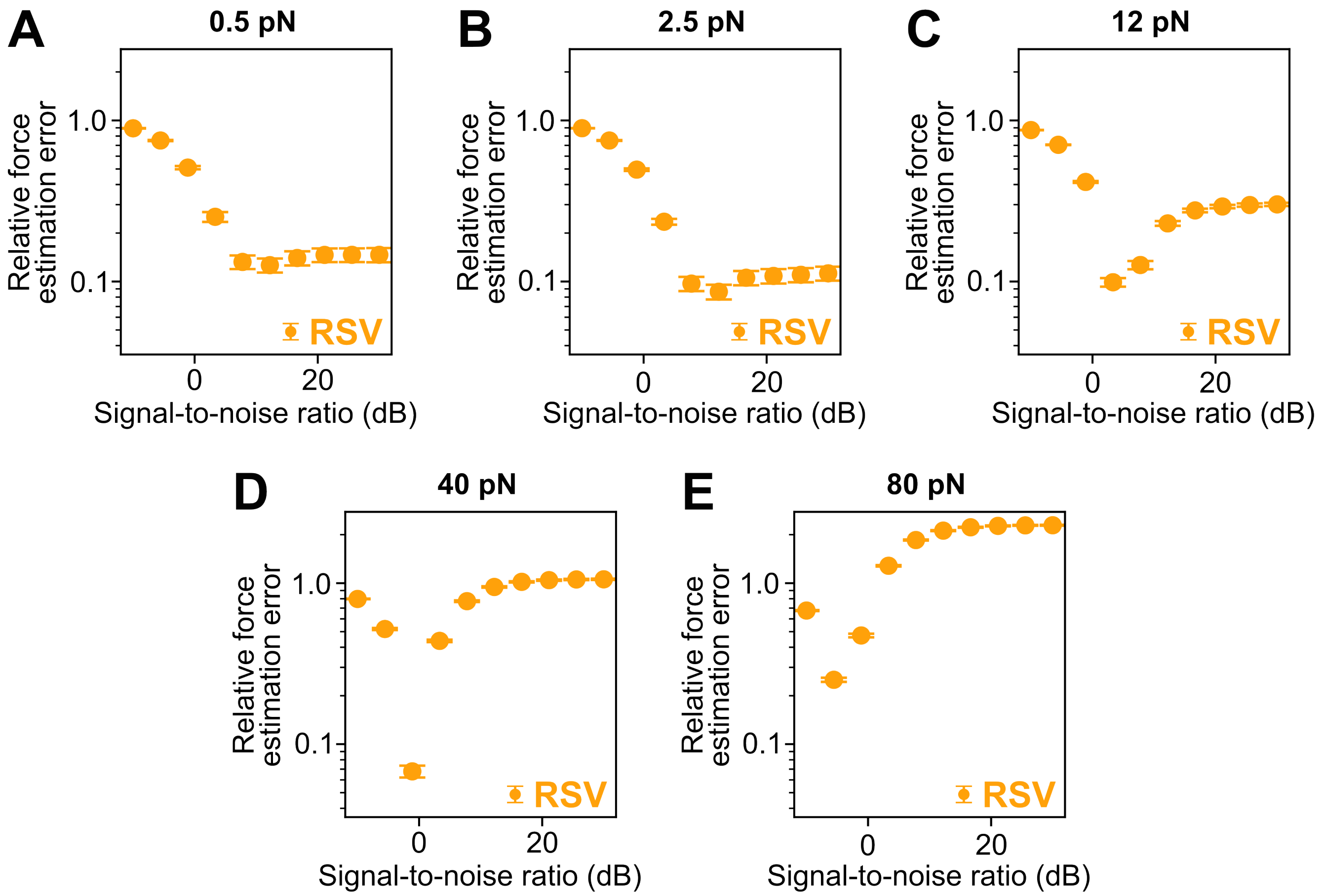


**Figure S17:** Effect of pink noise on the force estimation errors evaluated using simulated traces of 3 $\mu$m beads. Relative force estimation errors as a function of the signal-to-noise ratio for RSV at **(A)** 0.5 pN, **(B)** 2.5 pN, and **(C)** 12 pN, **(D)** 40 pN, and **(E)** 80 pN. The sampling frequency is 72 Hz.

#### $3\,\mu$m (M270) beads - Brownian noise

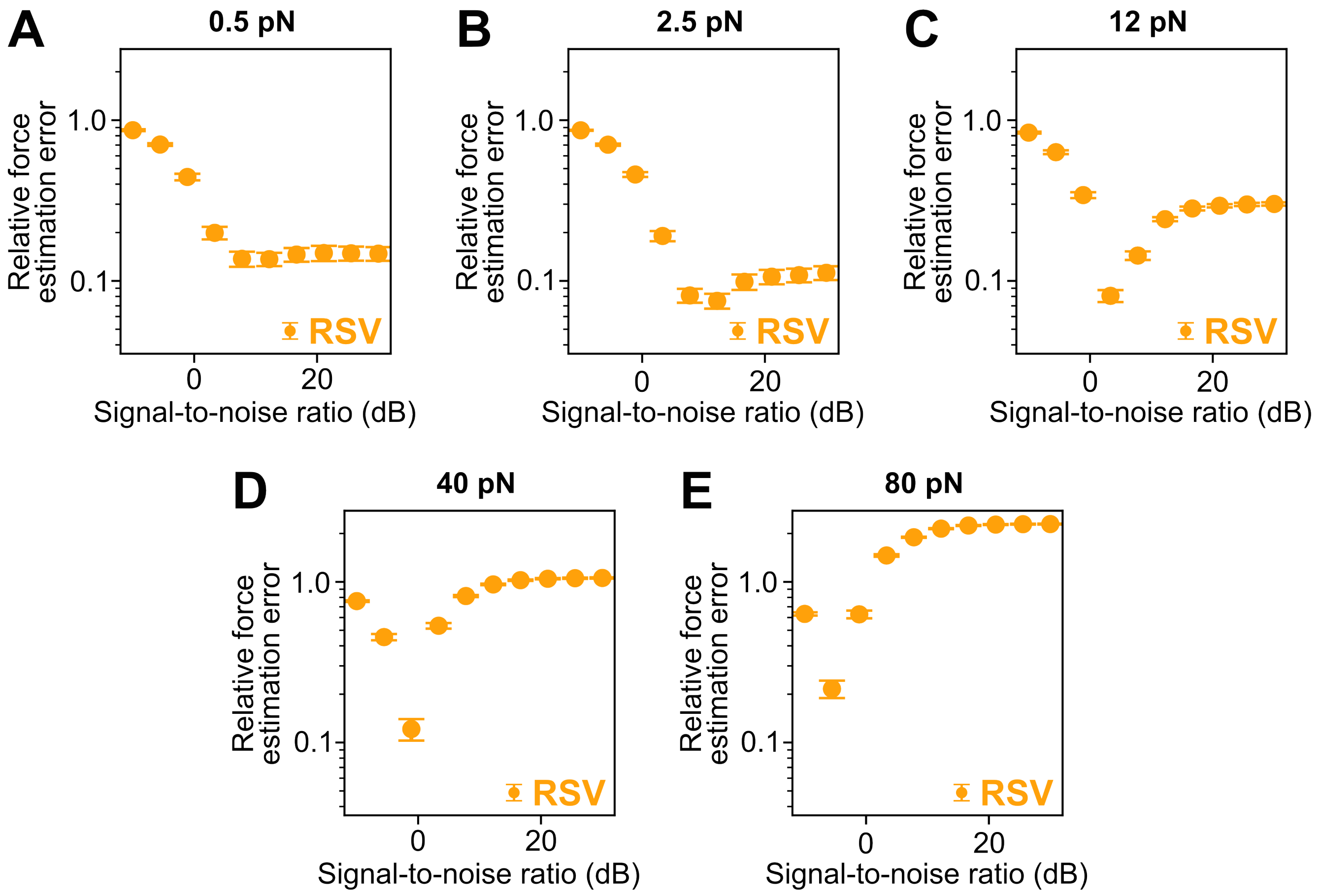


**Figure S18:** Effect of Brownian noise on the force estimation errors evaluated using simulated traces of $3\,\mu$m beads. Relative force estimation errors as a function of the signal-to-noise ratio for RSV at **(A)** 0.5 pN, **(B)** 2.5 pN, and **(C)** 12 pN, **(D)** 40 pN, and **(E)** 80 pN. The sampling frequency is 72 Hz.

3 $\mu$m (M270) beads - linear drift

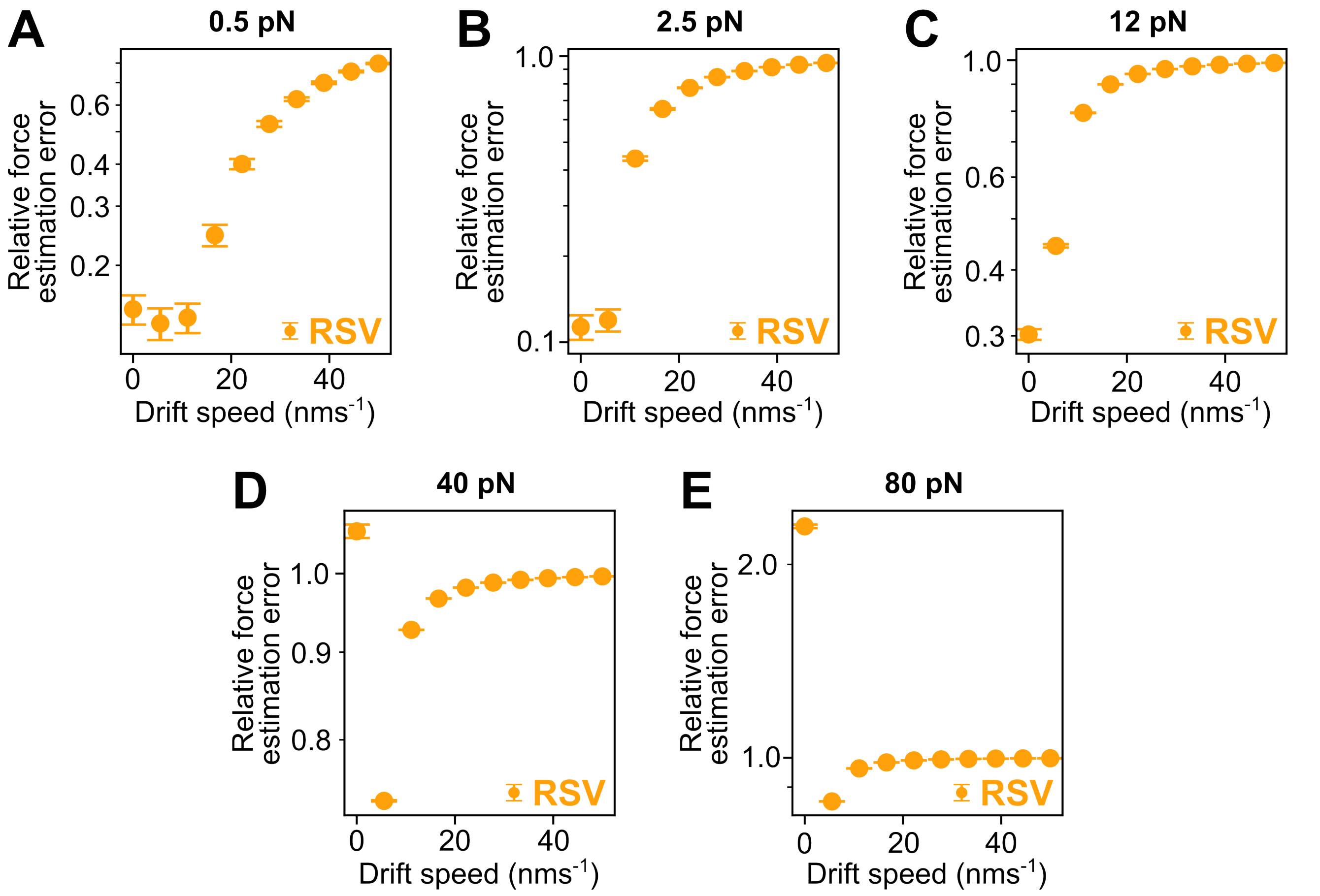


**Figure S19:** Effect of linear drift on the force estimation errors evaluated using simulated traces of 3 $\mu$m beads. Relative force estimation errors as a function of the drift speed for RSV at **(A)** 0.5 pN, **(B)** 2.5 pN, and **(C)** 12 pN, **(D)** 40 pN, and **(E)** 80 pN. The sampling frequency is 72 Hz.

#### $3\,\mu$m (M270) beads - non-linear drift

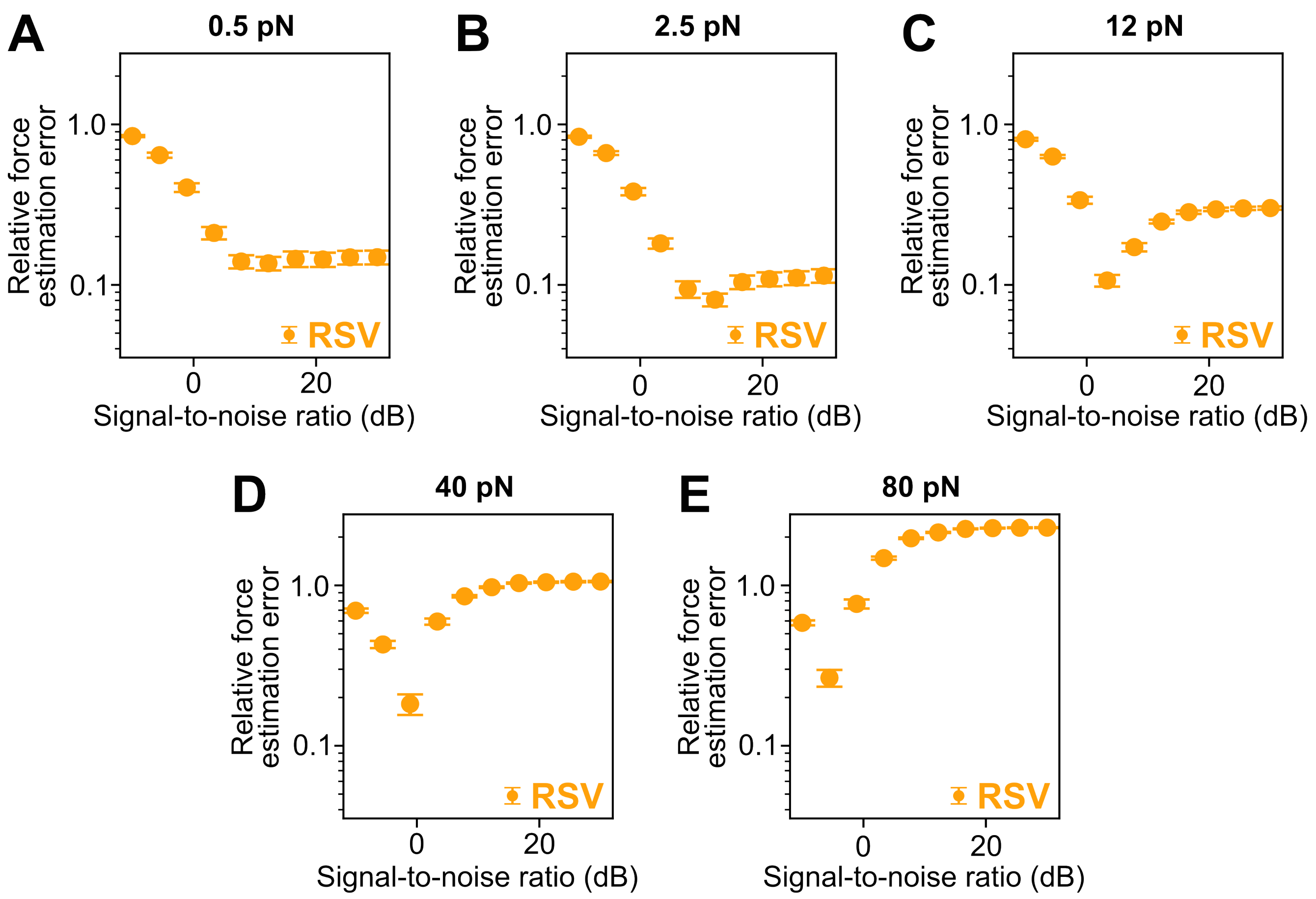


**Figure S20:** Effect of non-linear noise on the force estimation errors evaluated using simulated traces of $3\,\mu$m beads. Relative force estimation errors as a function of the signal-to-noise ratio for RSV at **(A)** 0.5 pN, **(B)** 2.5 pN, and **(C)** 12 pN, **(D)** 40 pN, and **(E)** 80 pN. The sampling frequency is 72 Hz.

### S8 Maximum Likelihood Estimation Fitting

To estimate the trap stiffness $\kappa$ and drag coefficient $\gamma$ of the measured or simulated bead trajectories, we first compute one of the methods, i.e PSD, AV, HV, using the collected trajectories. Then, we need to fit the corresponding analytical expression to the computed data. All the methods we have presented are quadratic in the positional data. Therefore, if we assume the uncertainties of the positional data are Gaussian distributed, the uncertainties of the computed data points follow a Gamma distribution which is given by

$$p(x; x^{\text{true}}, k) = \frac{1}{(k/x^{\text{true}}))^k \Gamma(k)} x^{k-1} e^{-kx/x^{\text{true}}}, \tag{16}$$

where $k$ is called the shape parameter and $\Gamma(k)$ is the Gamma function. This renders a least-squares fit non-optimal since least-squares method assumes a Gaussian distribution. Instead, we can use Maximum Likelihood Estimation (MLE) to fit the computed data to the correct distribution [7].

To perform MLE, we replace $x^{\text{true}}$ with $x^{\text{fit}}(\kappa, \gamma)$. Moreover, for PSD, $k$ is completely determined by the number of bins in its calculation. On the other hand, for AV and HV, $k$ depends on the dominant noise type within every bin[8]. Analytically exact values of $k$ exist for the power-law noises ($S(f) \propto f^{\alpha}$) for $\alpha \in [-4, 2]$ [9]. If the dominant noise type is out of this range, i.e., $\alpha \notin [-4, 2]$, $k$ can be approximated by [8]

$$k \approx \frac{1}{2}\left(\frac{N}{m} - 1\right), \tag{17}$$

where $N$ is the number of data points in the bead trajectory and $m$ is the bin size. Therefore, we eliminate the fitting parameter $k$ by using its exact or approximate value. Then, the only fitting parameters that are left are physical parameters, i.e. the trap stiffness $\kappa$ and drag coefficient $\gamma$. To perform the fitting, we construct the likelihood function as

$$L(\kappa, \gamma) = \prod_{i=1}^{N} p(x_i; \kappa, \gamma), \tag{18}$$

where $x_i$ denotes the $i^{\text{th}}$ data point. The best fit is at the point where this function takes its maximum value. To ensure numerical stability, we take its logarithm to get the log-likelihood funtion,

$$\ell(\kappa, \gamma) = \log L(\kappa, \gamma) = \sum_{i=1}^{N} \log p(x_i; \kappa, \gamma), \tag{19}$$

and maximize this function since its maximum is at the same point as the likelihood function. Therefore, the best fitting parameters are given by

$$(\hat{\kappa}, \hat{\gamma}) = \arg\max_{\kappa,\gamma} \ell(\kappa, \gamma). \tag{20}$$

To quantify the uncertainties of the fitting parameters, first, we evaluate the Hessian of the log-likelihood function at the best fitting parameters,

$$H(\hat{\kappa}, \hat{\gamma}) = \left.\begin{pmatrix} \frac{\partial^2 \ell}{\partial \kappa^2} & \frac{\partial^2 \ell}{\partial \kappa \partial \gamma} \\ \frac{\partial^2 \ell}{\partial \gamma \partial \kappa} & \frac{\partial^2 \ell}{\partial \gamma^2} \end{pmatrix}\right|_{(\hat{\kappa}, \hat{\gamma})}, \tag{21}$$

and obtain the covariance matrix using the identity given by

$$\text{Cov}(\hat{\kappa}, \hat{\gamma}) = 2H^{-1}(\hat{\kappa}, \hat{\gamma}). \tag{22}$$

Furthermore, we calculate the reduced chi-squared statistic ($\chi^2_\nu$) of the regression as

$$\chi^2_{N-2} = \frac{1}{N-2} \sum_{i=1}^{N} \frac{(x_i - x_i^{\text{fit}}(\hat{\kappa}, \hat{\gamma}))^2}{\sigma^2_{x_i}}, \tag{23}$$

where $\sigma_{x_i}$ is the uncertainty of $x_i$. Finally, we calculate the uncertainty of the fit parameters using the covariance matrix and the reduced chi-squared statistic by

$$(\sigma_{\hat{k}}, \sigma_{\hat{\theta}}) = \chi^2_{N-2} \sqrt{\text{diag}(\text{Cov}(\hat{\kappa}, \hat{\gamma}))}. \tag{24}$$

#### S9 Experimental Force Estimation Errors of M270 beads

The relative force estimation errors of the experimental traces, where a 21 kbp dsDNA is attached to M270 beads, are calculated as described in the main text using the PSD, AV, and HV methods. For all three methods, the errors decrease with increasing force with overall lower errors in the case of the HV method (Fig. S21).

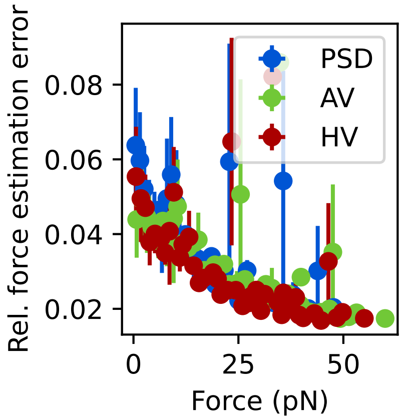


**Figure S21:** Relative force estimation errors as a function of force derived from experimental M270 data traces.

#### S10 Experimental Relative Bead-to-Bead Force Errors as a Function of the Force for MyOne Beads

To quantify the relative bead-to-bead force error, we calculate the bead-to-bead force errors from the relative force estimation errors presented in the main text (Fig. 5F) and fit the data (Fig. S22) with

$$\sigma_{\text{Force}} = \sigma_0 + \sigma_{rel} F, \tag{25}$$

where $\sigma_{\text{Force}}$ represents the bead-to-bead force errors in pN, $F$ (in pN) the force, and $\sigma_{rel}$ the unitless, relative bead-to-bead force error. The error $\sigma_0$ (in pN) accounts for intrinsic system errors. The derived values for $\sigma_{rel.}$ and $\sigma_0$ using the PSD, AV, and HV method are summarized in Table 1.

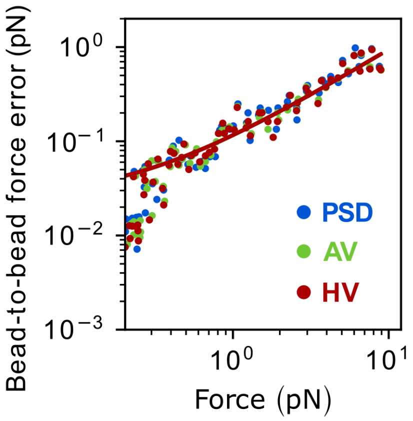


**Figure S22:** Experimental bead-to-bead force errors as a function of the force derived from experimental data traces using 21 kbp dsDNA and MyOne beads. For all three methods, the errors decrease with increasing force without preference for one type of method.

| | $\sigma_{rel.}$ | $\sigma_0$ (pN) | $r^2$ |
|---|---|---|---|
| PSD | $0.115 \pm 0.005$ | $0.005 \pm 0.011$ | 0.92 |
| AV | $0.110 \pm 0.004$ | $0.005 \pm 0.009$ | 0.94 |
| HV | $0.109 \pm 0.004$ | $0.004 \pm 0.009$ | 0.93 |

**Table 1:** Fitting values of the bead-to-bead force error analysis using Eq. 25 for the PSD, AV, and HV methods.
